# Inter-species comparison of the orientation algorithm directing larval chemotaxis in the genus *Drosophila*

**DOI:** 10.1101/2021.09.17.460740

**Authors:** Elie Fink, Matthieu Louis

## Abstract

Animals differ in their appearances and behaviors. While many genetic studies have addressed the origins of phenotypic differences between fly species, we are still lacking a quantitative assessment of the variability in the way different fly species behave. We tackled this question in one of the most robust behaviors displayed by Drosophila: chemotaxis. At the larval stage, *Drosophila melanogaster* navigate odor gradients by combining four sensorimotor routines in a multilayered algorithm: a modulation of the overall locomotor speed and turn rate; a bias in turning during down-gradient motion; a bias in turning toward the gradient; the local curl of trajectories toward the gradient (“weathervaning”). Using high-resolution tracking and behavioral quantification, we characterized the olfactory behavior of eight closely related species of the Drosophila group in response to 19 ecologically-relevant odors. Significant changes are observed in the receptive field of each species, which is consistent with the rapid evolution of the peripheral olfactory system. Our results reveal substantial inter-species variability in the algorithms directing larval chemotaxis. While the basic sensorimotor routines are shared, their parametric arrangements can vary dramatically across species. The present analysis sets the stage for deciphering the evolutionary relationships between the structure and function of neural circuits directing orientation behaviors in Drosophila.

## Introduction

Behavior enables animals to actively sense and adapt to their environment: it maximizes survival by allowing adaptive responses to external cues. The meaning of an environmental signal depends on the ecosystem in which an animal is immersed. While an odorant molecule might be predictive of food in a particular ecological niche, the same odorant molecule might have no association with food in a different ecological niche. As a result, sensory systems have evolved to be most sensitive to the stimuli an animal encounters and values in its natural environment (Laughlin 1981). Behavioral adaption, however, is not strictly limited to changes in the peripheral sensory system. It also emerges from changes in higher-order functions, such as the acquisition of social traits in feeding behavior (De Bono and Bargmann 1998) or the specialization of motor programs to cope with the properties of different physical environments (Karageorgi, Bräcker et al. 2017). Consistent with the idea that evolution can act by manipulating the properties of neural circuits, variations in the courtship rituals of drosophilids has been associated with changes in the functional logic of central circuits without major differences in the ability to detect pheromonal cues at the sensory periphery (Seeholzer, Seppo et al. 2018, Khallaf, Auer et al. 2020, Sato, Tanaka et al. 2020). While the genetic mechanisms underlying the functional diversification of neural circuits remain poorly understood, species-specific differences in courtship songs have been linked to differences in the genomic sequences of an ion channel widely expressed in the fly nervous system (Ding, Berrocal et al. 2016).

The extent to which behavioral adaptation is driven by changes at the sensory periphery versus higher-order brain functions remains elusive. To address this question, we studied the variability underlying chemotaxis in larvae of the Drosophila group. The reasons for this choice are twofold. First, olfactory behaviors are expected to display considerable variability across species (Bendesky and Bargmann 2011) given that odorant receptors are among the fastest evolving gene family (Sánchez-Gracia, Vieira et al. 2009). Second, larval chemotaxis involves a combination of stereotyped behaviors that can be quantified and classified computationally (Gomez-Marin and Louis 2012). Larval chemotaxis relies on a hierarchy of sensorimotor processes —routines— summarized in Figure 1. Together, these elementary routines form a behavioral algorithm. The basic algorithm directing larval chemotaxis is well-characterized in *Drosophila melanogaster*. The larva moves through peristaltic locomotion (Heckscher, Lockery et al. 2012). The propagation of each wave of muscle contraction can be viewed as a stride. Between two rounds of peristaltic contraction, the larva can adjust its direction of motion (Gomez-Marin and Louis 2014). Interruption of peristalsis produces stopping during which the larva is still able to sweep its head laterally (Lahiri, Shen et al. 2011) — a process called “head casting”. Stops are often followed by an abrupt change in orientation producing “turning”. From a macroscopic perspective, the larva alternates bouts of relatively straight motion with abrupt changes in reorientation (Figure 1A). In essence, this algorithm is analogous to bacterial chemotaxis where “runs” alternate with “turns” (also called “tumbles”) (Berg 2008), which explains why the same terminology has been applied to describe larval chemotaxis (Gomez-Marin and Louis 2012).

**Figure 1:**
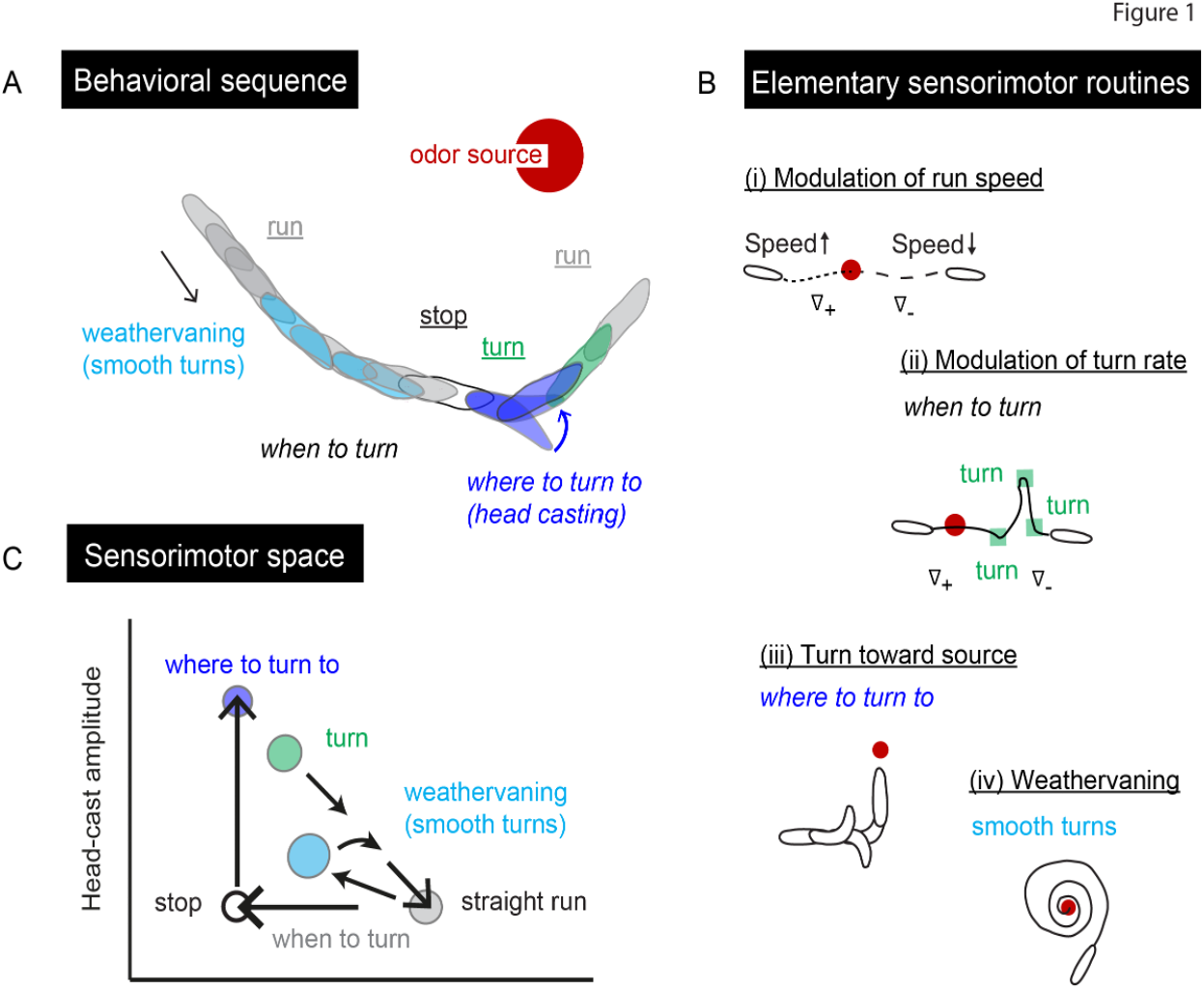
Conceptual framework of the reorientation algorithm directing larval chemotaxis toward attractive odors in *D. melanogaster*. **(A)** Fictive behavioral sequence of a larva moving in the vicinity of an odor source (red dot). Successive contours of the larva describe a run followed by a turn. The three fundamental behavioral states are: runs, stops and turns. Runs can be straight (gray) or they can curl toward/away from the source (light blue). Stops are usually associated with lateral head casts (dark blue) and followed by turning (green) when peristalsis resumes. **(B)** Four elementary sensorimotor routines underlying larval chemotaxis: (i) modulation of the run speed as a function of the bearing to the source. Larvae tend to accelerate upgradient and decelerate downgradient. (ii) Modulation of the turn rate, also referred to as *when-to-turn* routine. The probability of stopping increases downgradient and decreases upgradient. (iii) Bias of turning direction toward the odor gradient as a result of active sampling through head casts. This routine is also referred to as *where-to-turn-to* routine. (iv) Steering of the run direction toward the gradient through smooth turns in a process called *weathervaning*. **(C)** Transitions between the main behavioral states described in panel 1A in a space that is spanned by the run speed and head-cast amplitude. Straight runs correspond to high run speed and low head cast. Weathervaning corresponds to high run speed and medium turn rate. Stops are characterized by low run speed with or without head casts. During turning, both the run speed and head-cast amplitude are high. Panel 1C is adapted from the Figure 9 of (Gomez-Marin and Louis 2014).

When stimulated by a food odor gradient, larval chemotaxis involves four basic routines that convert sensation into the motor control directing runs and turns (Figure 1B): (i) The first sensorimotor routine modulates the run speed based on the heading with respect to the gradient’s direction. While larvae accelerate during upgradient motion, they decelerate during downgradient motion (Gershow, Berck et al. 2012, Schleyer, Reid et al. 2015). (ii) A hallmark of bacterial chemotaxis is the suppression of turning during runs oriented toward positive gradients. Larvae are capable of bidirectional modulation of their turn rate: positive gradients result in a decrease in turn rate while negative gradients result in an increase in turn rate (Gomez-Marin, Stephens et al. 2011). Here, we will refer to this sensorimotor routine as the ability to control the timing of stop-turns or the “*when-to-turn*” routine (Figure 1B). (iii) Overall, bacteria turn at random without a bias toward the direction of the source. Conversely, larvae are capable of biasing a majority of their turns toward the odor gradient (Gomez-Marin, Stephens et al. 2011). This process involves active sensing of the gradient through head casts — a routine we will refer to as “*where-to-turn-to*” (Figure 1B). (iv) Runs implemented by a larva are not perfectly straight: on average, runs tend to bend toward the direction of the local odor gradient. This process called “*weathervaning*” was first reported in the nematode *C. elegans* (Iino and Yoshida 2009). The sensorimotor routine underlying weathervaning features smooth turns without any interruption in peristalsis (Gomez-Marin and Louis 2014) (Figure 1B and 1C).

According to the framework described in Figure 1B, the behavioral state of a larva can be mapped onto a space described by two motor variables: the run speed and the amplitude of the head casts (Figure 1C). Rather than producing discontinuous jumps between discrete states, larval chemotaxis is thought to form a continuum in the transitions between behavioral states (Gomez-Marin and Louis 2014). Behavioral transitions are nonetheless highly stereotyped, which enables a reliable classification of behavioral modes both manually and computationally (Green, Burnet et al. 1983, Gomez-Marin, Stephens et al. 2011, Gershow, Berck et al. 2012, Schulze, Gomez-Marin et al. 2015). At the larval stage, virtually nothing is known about the odor preferences and chemotactic strategies of species other than *D. melanogaster*. Building on our conceptual framework for larval chemotaxis in *D. melanogaster* (Figure 1), we characterized and compared the sensorimotor algorithms used by larvae of 8 species when stimulated by a panel of 19 odorant molecules with ecological relevance. By applying computational techniques to correlate sensation with reorientation maneuvers, we performed a detailed inspection of the sensorimotor routines directing chemotaxis across species. We found that the 19 tested odors can be clustered into stimuli that elicit either conserved or variable responses. The screen reveals a large degree of variability in the use of sensorimotor routines, suggesting that the basic sensorimotor routines undergo different evolutionary constraints based on their necessity to mediate strong chemotaxis toward ecologically meaningful odorant molecules.

## Results

To determine how the algorithm directing larval chemotaxis has evolved during the course of speciation, we screened the behavior of larvae of a subset of closely related fly species. We selected eight representative species of the Drosophila group. The six species that belong to the melanogaster subgroup populate diverse regions of sub-Saharan Africa: *D. melanogaster*, *D. simulans*, *D. sechellia*, *D. erecta*, *D. santomea* and *D. yakuba* (Figure 2A 1-6). In spite of their striking anatomical resemblance, *D. melanogaster* and *D. simulans* are not sympatric: their geographical distributions differ within continental Africa (Lachaise, Cariou et al. 1988). Unlike *D. melanogaster*, *D. simulans* can thrive independently of human activities. Both *D. melanogaster* and *D. simulans* are generalists that feed on a variety of fermenting substrates. *D. melanogaster* feed on shrubs, figs, to name a few, with a particular attraction to the Marula fruit (Mansourian, Enjin et al. 2018). Although the exact ecological niche of *D. simulans* remains to be elucidated, it is associated with grass plants species (David, Lemeunier et al. 2007).

**Figure 2:**
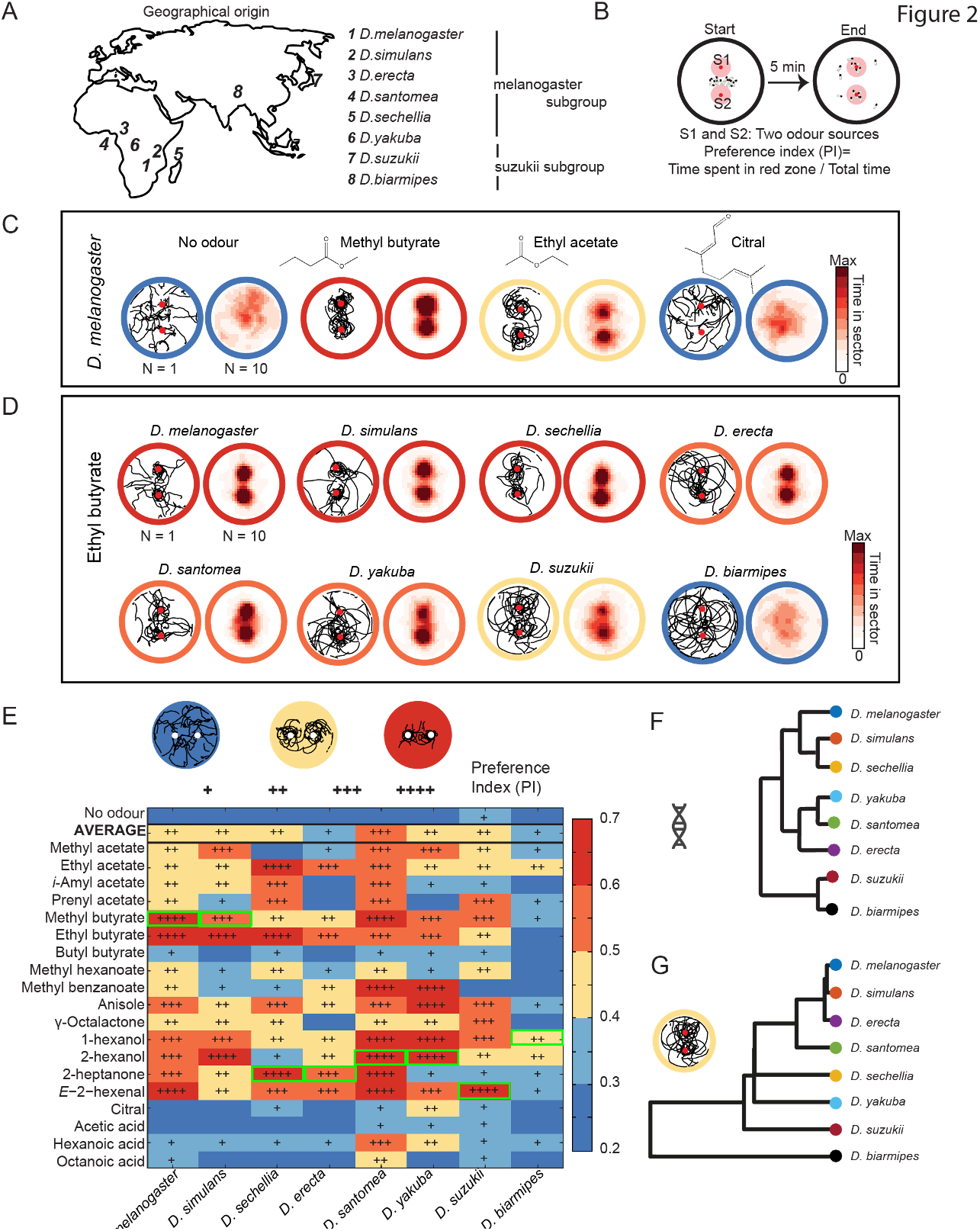
Assessing similarities and differences in odor preferences across *Drosophila* species. **(A)** Geographical origin of the species. **(B)** Schematic of the behavioral assay used in the odor screen. In each trial, 15 larvae of one species at a time are placed in the center of a 15-cm Petri dish in the absence (no-odor control) and presence of odor. The odor is loaded at two positions (P1 and P2) onto the Petri dish lid. The position of each larva is monitored for 5 minutes. The preference index (PI) is then calculated, defined as the proportion of time spent within the red area around the sources. (**C)** Four representative trials illustrating the no-odor condition and odors eliciting strong, medium, weak attraction in *D. melanogaster* larvae. The trajectories were obtained from 15 larvae in a single trial. **(D)** *Drosophila* species exhibiting different responses to the same odor stimulus, ethyl butyrate ranging from strong (dark red) to weak (blue) attraction. **(E)** Heatmap of the responses of each species to 19 odors, as well as the average response of one species across all odors. The level of attraction is quantified with the preference index (PI). **“-”** PI < 0.3; **“+”** 0.3 ≤ PI < 0.4; **“++”** 0.4 ≤ PI < 0.5; **“+++”** 0.5 ≤ PI < 0.6; **“++++”** 0.6 ≤ PI < 0.7. The green squares indicate the odors forming the high-olfactory-preference (HOP) set for each species. The behavioral analysis will focus on the set of HOP odors in Figures 3, 4 and 5. Number of trials per experimental condition (species — odor): 10. **(F)** Consensus phylogenetic tree computed based on genomic information of the 8 species. Dendrogram adapted from (Seetharam and Stuart 2013). **(G)** Dendrogram of the olfactory behaviors computed based on a hierarchical cluster of the average response of each species measured in the response space (for details, see Materials and methods).

In contrast with *D. melanogaster* and *D. simulans*, *D. sechellia* and *D. erecta* are classified as “specialists” with respect to their feeding preferences (David, Lemeunier et al. 2007). *D. sechellia* is insular, endemic to the Seychelles islands where it reproduces and feeds on the Morinda fruit that is toxic to most other fly species. While *D. sechellia* displays multiple traits of behavioral specialization at the adult stage (Dekker, Ibba et al. 2006, Matsuo, Sugaya et al. 2007, Auer, Khallaf et al. 2020), little is known about the chemosensory behavior of this species at the larval stage. *D. erecta* is a seasonal specialist that feeds on the pandanus tree when present for several months. Although *D. yakuba* is evolutionarily closely related to *D. santomea*, the geographical distributions of these two species are strikingly distinct. *D. yakuba* is among the most abundant flies among the African endemic species: it is a domestic generalist that is widespread in sub-Saharan tropical Africa (David, Lemeunier et al. 2007) and feeds on a broad range of plants and fruits (Lachaise, Cariou et al. 1988). By contrast, *D. santomea* is a mountainous forest insular species endemic to the São Tomé and Príncipe Island. *D. santomea* can live at higher altitudes than *D. yakuba* and it feeds on fallen figs (Cariou, Silvain et al. 2001).

Two species represent the Asian suzukii subgroup: *D. suzukii* and *D. biarmipes* (Figure 2A). In recent years, *D. suzukii* has drawn a lot of attention due to the damages it inflicts on crops.

While this species was first described in Japan in 1931, its geographical distribution quickly expanded from Eastern to Southeastern Asia. *D. suzukii* has now become ubiquitous in Europe and Northern America (Hauser 2011). Due to the existence of an unusually large ovipositor to lay eggs in fresh fruits, *D. suzukii* — also known as the spotted wing drosophila or cherry vinegar fly — is considered as a global fruit pest that feeds on soft-skinned ripening fruit crops such as cherries, grapes and berries. Accordingly, *D. suzukii* demonstrates specialized olfactory and egg-laying behavioral traits at the adult stage (Karageorgi, Bräcker et al. 2017). Consistent with the observation that this species feeds and reproduces on fresh fruits, larvae of *D. suzukii* are capable of burrowing in harder substrates than *D. melanogaster* (Kim, Alvarez et al. 2017). In the present study, we used *D. biarmipes* as the sibling species of *D. suzukii*. Endowed with a regular ovipositor, *D. biarmipes* is unable to reproduce on fresh fruits: it feeds on soft fermenting fruits. Multiple differences in chemosensory and mechanosensory behaviors have been reported between *D. suzukii* and *D. biarmipes* (Karageorgi, Bräcker et al. 2017, Dweck, Talross et al. 2021).

To compare larval chemotaxis across species, we tracked the locomotor behavior of larvae in controlled odor gradients. A set of 19 odorant molecules (Figure 2E) was selected based on the biosynthesis of the molecules in native or introduced plant species reported as breeding sites in the fly’s natural environments (Lachaise, Cariou et al. 1988). The 19 odorant molecules included esters, alcohols, a ketone, aldehydes, an aromatic, a terpene, organic acids and compounds of varying chain lengths present in the headspace of fruits (Wick, Yamanishi et al. 1969, Jordán, Tandon et al. 2001, Grison-Pigé, Hossaert-McKey et al. 2002, Pino, Mesa et al. 2005, Dweck, Ebrahim et al. 2013, Kim, Kim et al. 2013, Linz, Baschwitz et al. 2013, Papa, Maggi et al. 2014). Odorants were tested at dilutions of 1:100 (0.01) or 1:200 (0.005) (see Materials and methods), which correspond to conditions used in several previous studies (De Bruyne, Foster et al. 2001, Hallem, Ho et al. 2004, Larsson, Domingos et al. 2004, Wilson, Turner et al. 2004, Kreher, Kwon et al. 2005, Kreher, Mathew et al. 2008). We note however that the final concentrations experienced by larvae in this assay might be slightly different from previous work given the differences in size of the behavioral assays and the odor volume applied.

The final subset of 19 odorant molecules was selected out of a larger group of odorant chemicals that were pre-screened on *D. melanogaster* larvae to assess whether they led to any olfactory response at all. This resulted in the exclusion of multiple odor candidates, particularly terpenes, which did not elicit chemotaxis behavior in *D. melanogaster.* Based on a pre-screen with anosmic *Orco-*null mutant larvae, we ensured that any odor dilution used in the screen did not elicit odor responses in anosmic *D. melanogaster* larvae. This measure was taken to avoid potential activation of other chemosensory receptors, such as gustatory receptors or ionotropic receptors, due to high concentrations, which would interfere with the behavior directed by airborne olfactory cues. A total of 152 species-odor combinations were tested (Figure 2E). As a control, we characterized the behavior of each species to the unstimulated (no-odor) condition.

The throughput of the screen was increased by testing the behavior of 15 larvae simultaneously in a group assay featuring 2 odor sources (Figure 2B). Under these conditions, the group of larvae tended to split equally between the two sources, which limited inter-individual collisions. Within a single species, *D. melanogaster*, responses ranged from strongly attractive to weakly attractive for different odors (Figure 2C). The attraction level to the source was quantified by a preference index (PI), which measures the average fraction of time larvae spent in a 1-cm radius zone surrounding the odor source (Figure 2B). The time courses of the distance to the odor source are presented in Supplementary Fig. 1.

We observed that a given odor could elicit responses ranging from strongly attractive to unattractive in all 8 species (Figure 2D). Ethyl butyrate, for instance, is a volatile ester widely found in fruits and it elicits both physiological and attractive behavioral responses in *D. melanogaster* adults and larvae (Stensmyr, Giordano et al. 2003, Kreher, Mathew et al. 2008, Asahina, Louis et al. 2009, Revadi, Vitagliano et al. 2015). This odor was clearly attractive to all species, except *D. biarmipes* (Figure 2D, bottom right). Conversely, some odors such as hexanoic acid and octanoic acid were only attractive to one or two species (Figure 2E). A summary of all PI results is compiled in Supplementary Fig. 2. We classified the 19 odors into 3 groups based on the overall level of attraction they elicited across the eight tested species (Supplementary Fig. 2).

Overall, each odor of the stimulus collection elicited significant attraction in at least one species (Figure 2E and Supplementary Fig. 2A). Every species responded to at least one odor, including *D. biarmipes*, which showed significant chemotaxis to ethyl acetate, 1-hexanol and 2-hexanol. A visual inspection of the trajectories suggested differences in chemotactic strategies between species. *D. biarmipes* was characterized by unusually circular trajectories (Figure 2D: *D. biarmipes* - ethyl butyrate), whereas the trajectories drawn by, for example, *D. melanogaster* to an odor of equivalent attraction level were more contorted and devoid of long uninterrupted circular segments (Figure 2C: *D. melanogaster* - citral). Interestingly, *D. santomea* demonstrated strong attraction to the largest number of tested odors (Figure 2E). Overall, this species tended to stay closest to the odor source for the duration of the trial (Supplementary Fig. 1).

Given the large number (152) of odor-species pairs, we focused our algorithmic analysis of chemotaxis on a subset of odors. To this end, we determined the high-olfactory-preference (HOP) odor that elicited the strongest PI in each species. The HOP odors are highlighted by a green square in Figure 2E. This subset of odors was used to standardize our analysis of the chemotaxis algorithms in Figures 4-6. In the case of *D. simulans*, we selected the second most attractive odor to use the same HOP condition as *D. melanogaster* and facilitate comparisons between the two species.

While on the one hand each of the eight species was characterized by a unique response profile to 19 tested odorant stimuli, the odors could produce homogeneous or heterogenous responses across species (Figure 3A). On average, ethyl butyrate was strongly attractive to the majority of the eight species (Figure 3A) with a high average PI taken across all species (Supplementary Fig. 2). Methyl benzoate produced heterogeneity in chemotactic responses across species, some showing strong attraction (e.g., *D. melanogaster*) and others only very weak attraction (e.g., *D. suzukii*) (Figure 3A), resulting overall in a medial average PI taken across all species (Supplementary Fig. 1). At the other end of the spectrum, we found that odors such as acetic acid fell into the category of stimuli that, on average, elicited a weak chemotactic response in all species (Figure 3A) and thus, a low average PI taken across all species (Supplementary Fig. 2).

**Figure 3:**
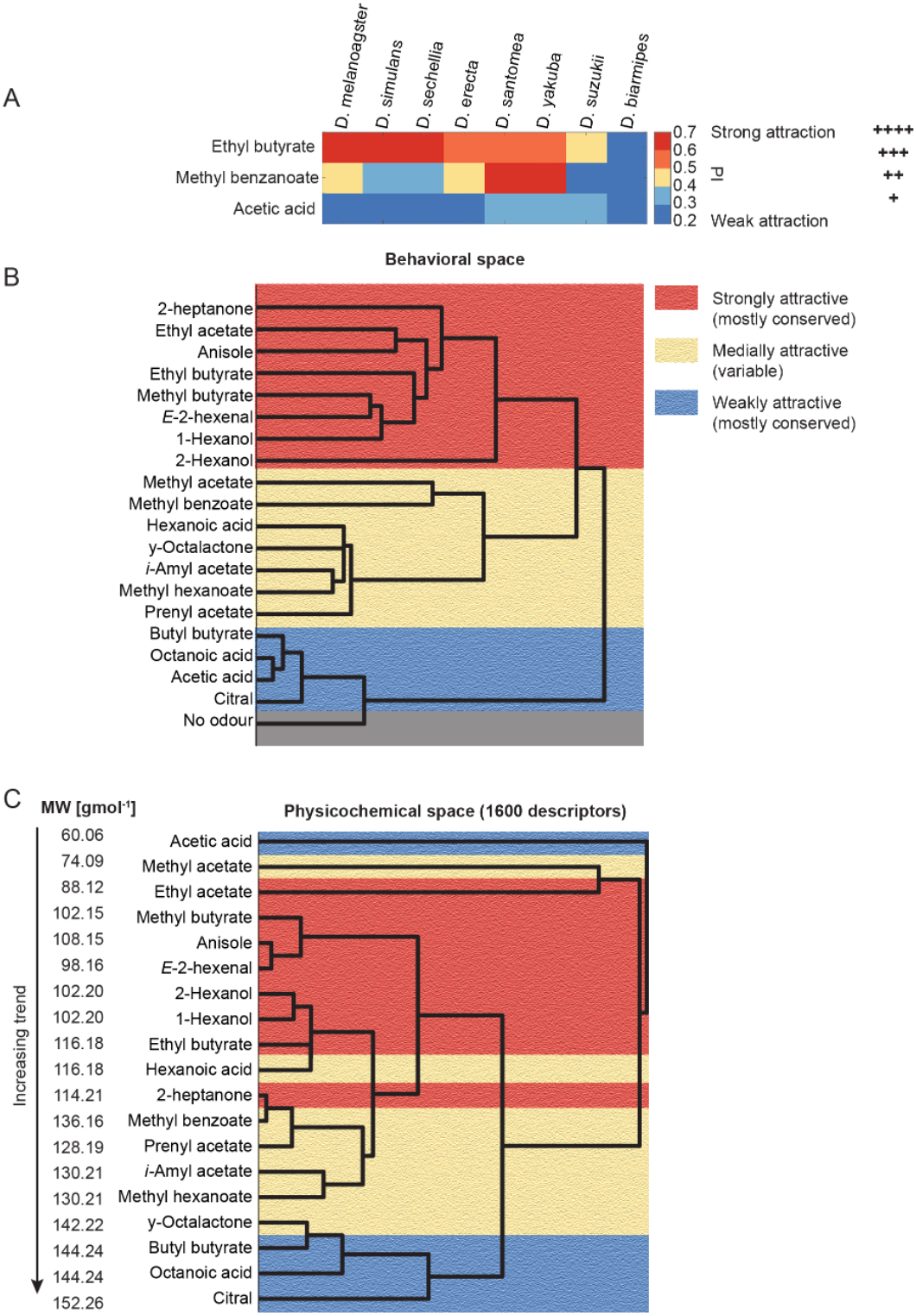
Variability and conservation of olfactory responses across species and across chemical structures. **(A)** Three conditions exemplifying how an odor can elicit conserved attraction in most species (ethyl butyrate), weak attraction (acetic acid) and variable levels of attraction across species (methyl benzoate). **(B)** Hierarchical cluster analysis based on the Euclidean distance measured between the preference indices (PI) elicited by the different odors for each species. **(C)** Hierarchical cluster analysis using the Euclidean distances between the physicochemical profiles of the odors (1600+ physicochemical descriptors reduced with a PCA, for details see Materials and methods).

Using the average PI of each odor, we constructed a similarity tree of the odors in the behavioral space by performing a hierarchical cluster analysis using the Euclidean distances between the odors based on the PI of each odor for each species as variable (Figure 3B).

The similarity tree allowed us to evaluate the closeness between the odors measured from the behavioral responses they elicited in individual species. Based on the structure of this tree, we defined three categories of odors (for detail, see Methods section): those that elicited strong attraction (red), variable and medial attraction (yellow), and weak attraction (blue) taken across all species (Figure 3B). The fact that the medially attractive odors tended to be associated with variable and heterogeneous responses across species could be seen by contrasting the attraction profiles of the strongly-attractive ethyl acetate and the medially- attractive prenyl acetate in Supplementary Fig. 2. Except for 2-hexanol and 2-heptanone, strongly attractive odors tended to produce inter-species homogeneity.

In analogy to the similarity tree in the behavioral space (Figure 3B), we constructed a similarity tree in the physicochemical space of the odors (Figure 3C). For this, we applied a principal component analysis (PCA) to the high-dimensional space of the odors described as 1600+ physicochemical descriptors. In the new reference system of the PCA, the coordinate axes correspond to principal components (PCs) that represent linear combinations of the molecular descriptors. In this space, we performed a hierarchical cluster analysis using the Euclidean distances between the odors. We found that the molecular weight descriptor greatly contributed to the variance along the first principal component (PC1), which is reflected in the distribution of the odors in the new PC space. Thus, the similarity tree shown in Figure 3C depicts the closeness between the odors primarily based on their molecular weight. According to this representation, the smallest molecules (e.g., acetic acid and methyl acetate) are furthest away from the largest molecules (e.g., octanoic acid and citral) in the similarity tree.

As a trend, we observed that odors eliciting strong attraction were predominantly smaller molecules with lower molecular weights, while odors eliciting weak attraction were predominantly larger molecules with higher molecular weights. A notable exception to this trend was acetic acid, which despite being the smallest molecule in the collection of stimuli, was very weakly attractive. This result might be explained by the fact that acetic acid is overall deleterious to *D. melanogaster* at the larval stage (Devineni, Sun et al. 2019). Medially attractive odors and odors with variable attraction levels across species fell between these two categories. Conserved odor responses, however, did span across chemical functional groups: for instance, the aldehyde *E*-2-hexenal and the alcohol 1-hexanol produced strong attractive responses with similar strengths.

*Drosophila* larvae navigate natural environments through an orientation algorithm that converts the detection of sensory stimuli into the control of motor programs underlying locomotion. Previous work has described that *Drosophila melanogaster* larvae orient through the sequential organization of stereotypical active-sensing behaviors described as runs, lateral head casts and turns (Gomez-Marin, Stephens et al. 2011, Gershow, Berck et al. 2012). As we found that these three basic motor programs were common to the eight tested species, we adopted the run speed and the turn rate (Figure 4A) as the principal sensorimotor variables controlling orientation responses. The turn rate quantifies maneuvers that involve stopping and abrupt changes in orientation. This process does not include smooth turns also known as *weathervaning* (Gomez-Marin and Louis 2014). Using the turn rate and speed to characterize the response elicited by the HOP odors, stark differences were observed between the eight species (Figure 4B). Both during odor-stimulated and unstimulated foraging, the eight species separated into two groups evaluated by k-means clustering (Figure 4B and Supplementary Fig. 3). This clustering resulted from differences in the absolute values of the turn rate and speed of each species. Using a linear discriminant analysis, we established that locomotor strategies significantly differed between the 8 species (Figure 4B).

**Figure 4:**
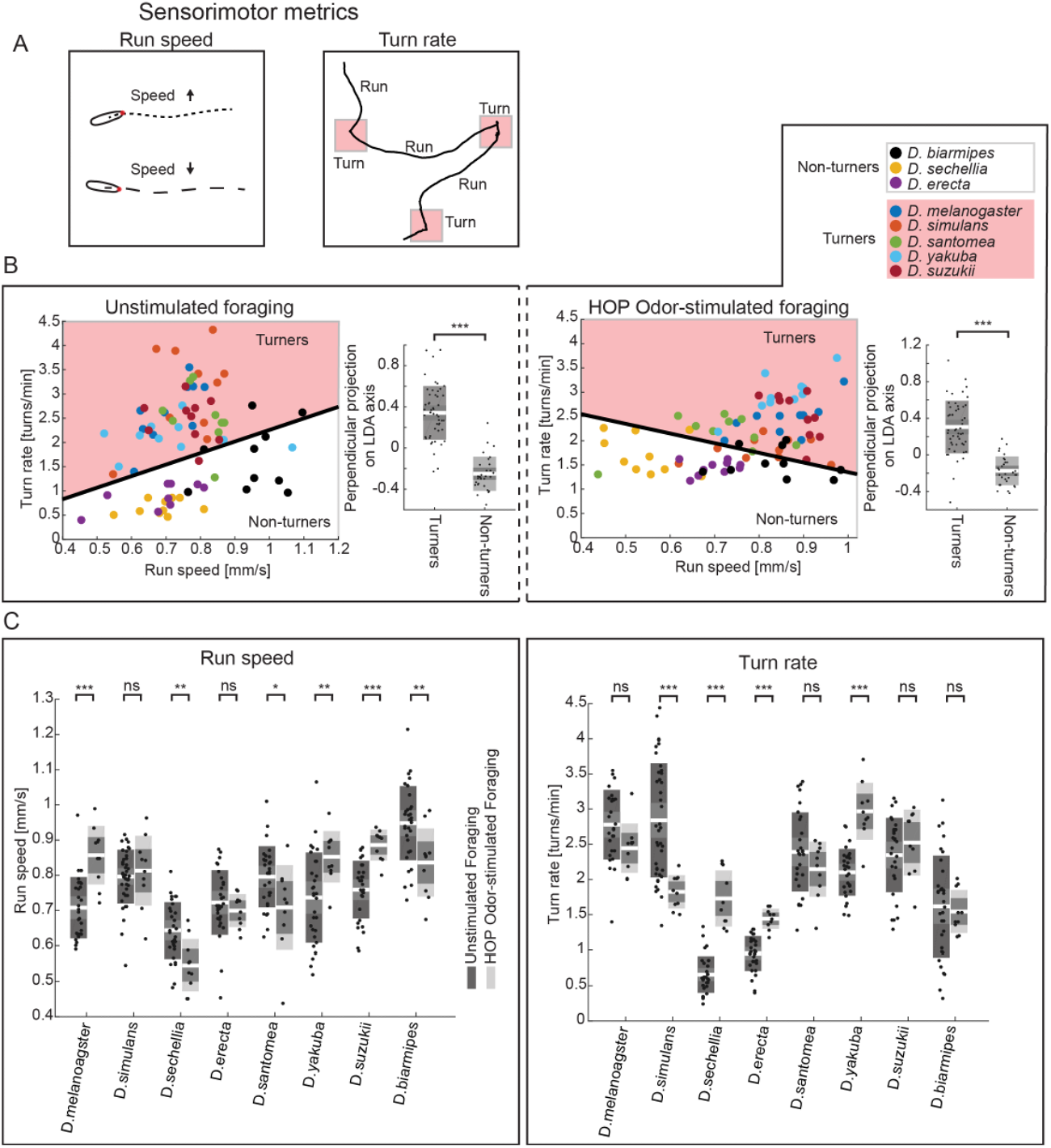
Two sensorimotor variables differentiate the control of orientation responses during foraging and chemotaxis across species. **(A)** Two sensorimotor metrics, run speed (speed of the midpoint) and turn rate, represent the main variables underlying behavioral responses during foraging and chemotaxis across species. **(B)** Linear decomposition analysis (LDA) of the sensorimotor metrics, run speed and turn rate, associated with unstimulated (left panel) and HOP odor-stimulated behaviors (right panel). The results of the LDA suggest the existence of two clusters of species separated by their turn rates: the “turners” and “non-turners”. The box plots show a perpendicular projection of the behavioral data onto the LDA axis (thick black line in the scatter plot of the run speed and turn rate). Differences between the projected means of the turners and non-turners were tested by a two-sample *t*-Test (***p≤0.001). Individual dots correspond to the average speeds and turn rates calculated during a particular trial. Number of trials: 10. **(C)** The average turn rate and run speed during HOP-stimulated behaviors are distinctly modulated up, down or not at all in each species compared to the unstimulated foraging. Differences between the means of the HOP-stimulated and the unstimulated behaviors were tested by a two-sample *t*-Test (*p≤0.05, **p≤0.01, ***p≤0.001, ns: not significant *p*>0.05). Number of trials per experimental condition (species — odor): 10. In the boxplot, the mean (white line) is surrounded by a gray area that corresponds to 1.96 SEM (95% confidence interval), and a colored interval that corresponds to 1 SD.

Next, we considered how each species modulates its speed and turn rate in response to the HOP stimulation relative to the unstimulated condition. As seen in Figure 4C, these two metrics were up- or down-regulated in a species-specific manner. In a given species, the nature of the modulation induced by the odor could not be predicted based on the fact that this species was classified as a turner or a non-turner. Belonging to either group was not predictive of differences in the overall level of chemotaxis. For instance, being a non-turner did not mean that a species was not capable of chemotaxis. Although *D. sechellia* was classified as a non-turner (Figure 4B, top insert), its low turn rate increased upon HOP stimulation compared to the unstimulated condition. By contrast, *D. santomea* did not modulate its high turn rate upon HOP stimulation even though it belonged to the turner group (Figure 4C).

In Figures 5 and 6, we inspected how the sensory experience of an animal can further modulate its run speed as well as other aspects of the navigation behavior, specifically *when-to-turn*, *where-to-turn-to* and *weathervaning* (Gomez-Marin and Louis 2014). We started by considering the ability of a larva to modulate its speed of locomotion based on information extracted from the odor gradient (Figure 5Ai). First, most species underwent a change in speed upon HOP stimulation (Supplementary Fig. 4A). While *D. melanogaster*, *D. yakuba* and *D. Suzukii* speeded up, *D. sechellia* and *D. santomea* slowed down. Whether the detection of the odor produced an overall increase or decrease in speed was a feature that tended to be conserved across all tested odors, except for *D. simulans*, *D. erecta* and *D. biarmipes* that showed considerable inter-odor variability (Supplementary Fig. 4B).

**Figure 5:**
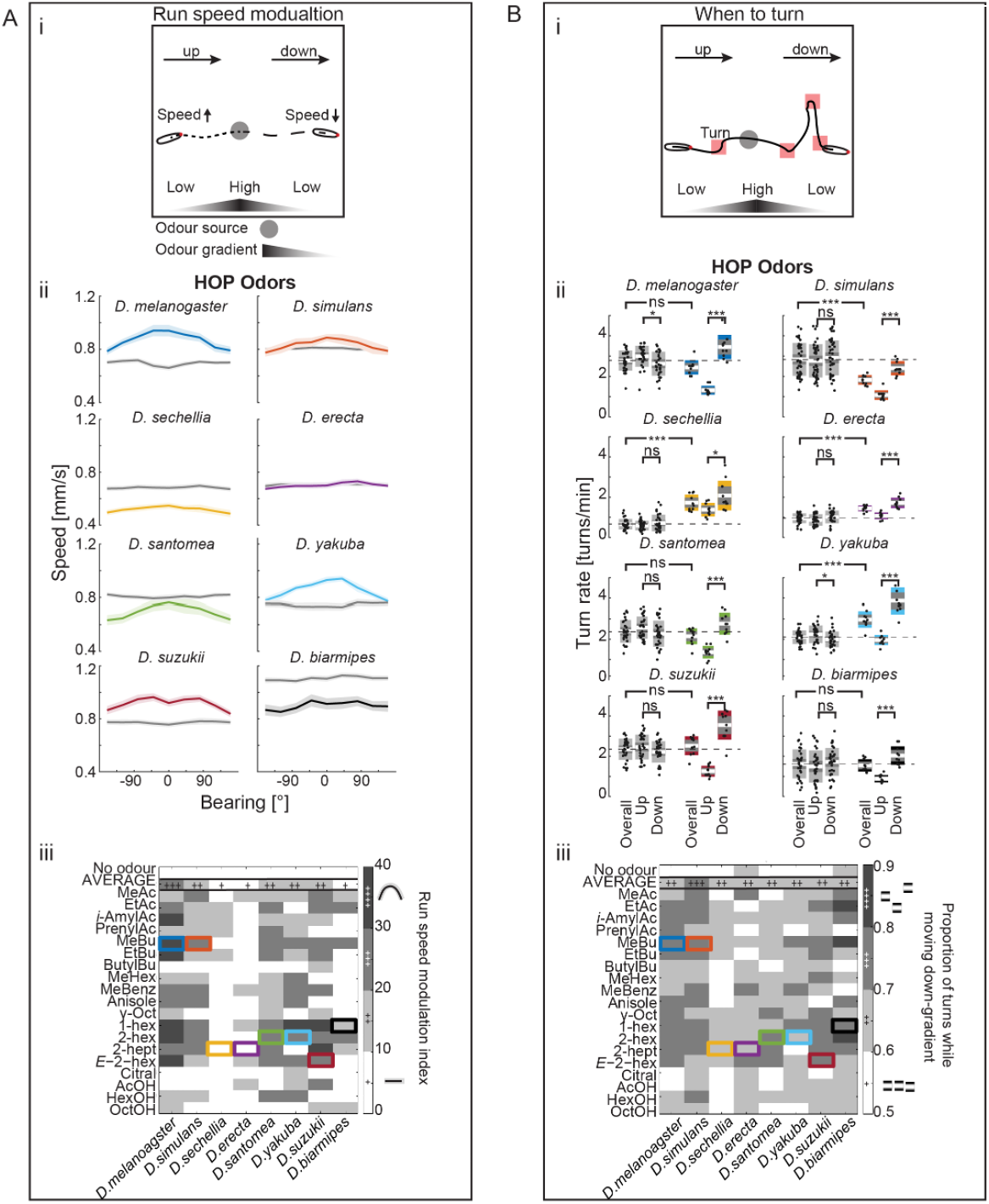
Modulation of the speed and turn rate by the animal’s orientation in the odor gradient. **(Ai)** The run speed is modulated as a function of the animal’s orientation in the odor gradient during chemotaxis. **(Aii)** The high-olfactory-preference (HOP) set of odors results in a distinct modulation of the run speed by each species as a function of the bearing to the odor gradient. Thick lines represent the mean speeds surrounded by their SEM displayed as shaded areas. **(Aiii)** Heat map of the run speed modulation index for all species and all tested odors. The value for the HOP odor of each species is demarked by an open boxed colored according to the convention used in Aii. **(Bi)** The turn rate is modulated as a function of the animal’s orientation in the odor gradient during chemotaxis, resulting in an increase or decrease of the turn rate depending on whether the larva is moving towards or away from the odor (*when-to- turn* modulation). **(Bii)** The HOP set of odors results in distinct modulation of the *when-to-turn* behavior by each species as a function of the bearing to the odor gradient. Differences of overall turn rates of odors with unstimulated condition, as well as differences of turn rates up and down for each condition were tested by a two-sample *t*-Test (*p≤0.05, **p≤0.01, ***p≤0.001, ns: not significant *p*>0.05). In the boxplot, the mean (white line) is surrounded by a gray area that corresponds to 1.96 SEM (95% confidence interval), and a colored interval that corresponds to 1 SD. **(Biii)** Heat map of the proportion of turns while moving down-gradient for all species and all tested odors. The value for the HOP odor of each species is demarked by an open-colored box. Number of trials per experimental condition (species — odor): 10.

**Figure 6:**
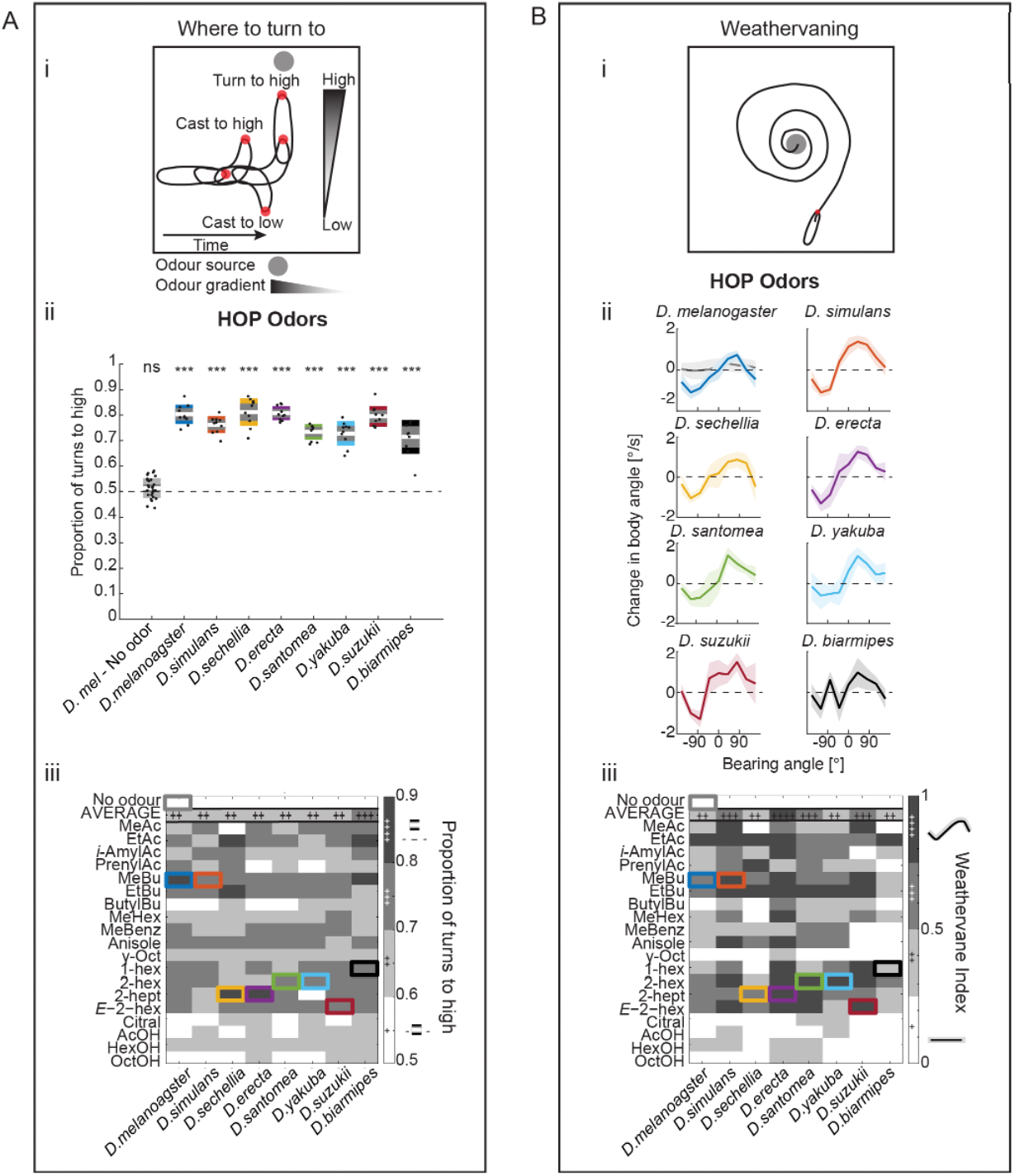
Modulation of the turning maneuvers by the animal’s orientation in the odor gradient. **(Ai)** The direction of a turn depends on the animal’s orientation in the odor gradient during chemotaxis. **(Aii)** The high-olfactory-preference (HOP) set of odors for each species results in consistently higher proportions of turns towards the odor gradient compared to the absence of an odor. Differences of turning proportions with chance were tested by a one-sample one-sided *t*-Test (\**p*≤0.05, \*\**p*≤0.01, \*\*\**p*≤0.001, ns: not significant *p*>0.05). In the boxplot, the mean (white line) is surrounded by a gray area that corresponds to 1.96 SEM (95% confidence interval), and a colored interval that corresponds to 1 SD. **(Aiii)** Heat map of the proportions of turns to high (turns toward the odor source) for all species and all tested odors. The value for the HOP odor of each species is demarked by an open colored boxed. **(Bi)** The turning behavior of an animal can also be achieved in continuous smaller reorientations as a function of the animal’s orientation (bearing) in the odor gradient during chemotaxis (weathervaning). **(Bii)** The HOP set of odors results in weathervaning behavior by each species as a function of the HOP gradient. Thick lines represent the mean reorientation rates surrounded by their SEM displayed as shaded areas. **(Biii)** Heat map of the weathervaning index for all species and all tested odors. The weathervaning index ranges between 0 (absence of correlation between the reorientation rate and the bearing) and 1 (perfect correlation) (see Materials and methods). The value for the HOP odor of each species is demarked by an open colored boxed. Number of trials per experimental condition (species — odor): 10.

In addition to an overall change in speed, larvae could adjust their run speed relative to their movement with respect to the odor gradient. This effect has been reported in *D. melanogaster* (Schleyer, Reid et al. 2015), which accelerates while moving up-gradient and decelerates while moving down-gradient (Figure 5Aii, top left). While some species displayed a strong increase in the run speed during movements towards the HOP source (e.g., *D. yakuba*), other species did not modulate the run speed at all (e.g., *D. erecta*). As will become clear in the rest of the analysis (Figure 6), a lack of modulation of the run speed in response to the local gradient did not imply that a species was unable to detect changes in odor concentration.

Next, we inspected species-specific differences in the *when-to-turn* routine, which corresponds to the ability of a larva to turn with a higher frequency during down-gradient motion and to turn less frequently during up-gradient motion (Figure 5Bi). While the baseline turn rate was unique to each species, resulting in the separation of species into turners and non-turners (Figure 4B), the sensory-dependent modulation of the turn rate was a feature common to all species upon HOP stimulation (Figure 5Bii). This result could be extended to the majority of odors tested in the study (Figure 5Biii), but a counter-example is highlighted in Figure 7. Thus, *when-to-turn* is a primary sensorimotor routine that is conserved across all species.

**Figure 7:**
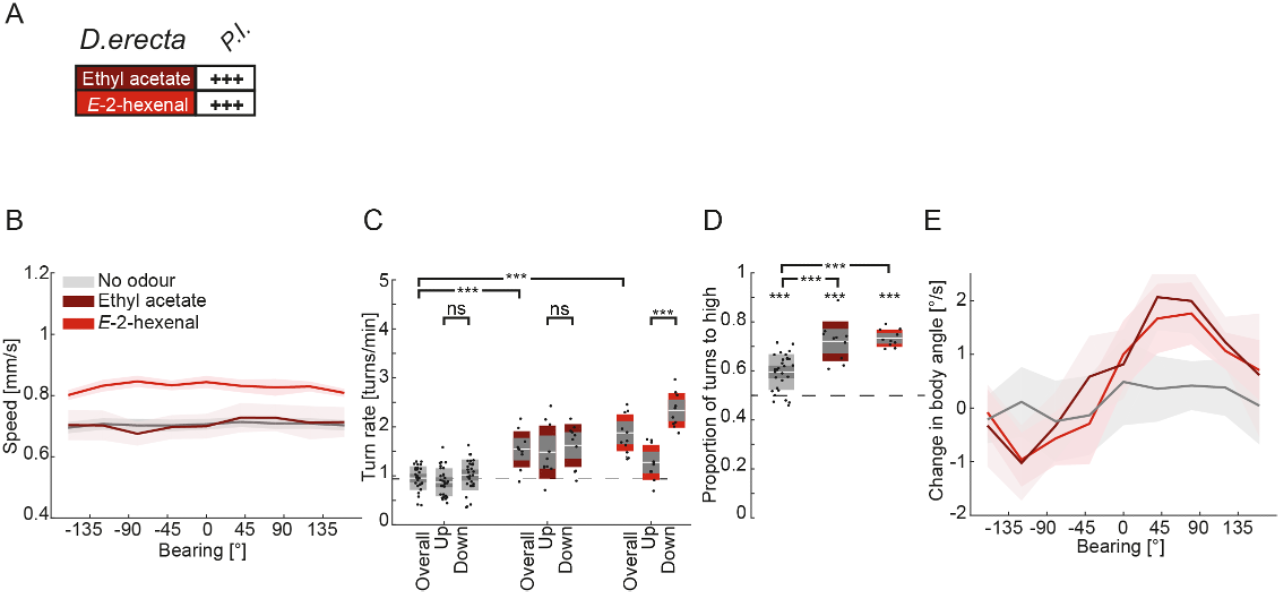
Intra-species variability: similar attraction levels to different odors can arise from different sensorimotor control in one species. **(A)** Ethyl acetate and *E*-2-hexenal elicit high preference indices (PI) indicative of strongly attractive responses in *D. erecta*. **(B)** *D. erecta* displays different run speeds in the presence of ethyl acetate and *E*-2-hexenal. The run speed remains the same as in the no-odor control for ethyl acetate, while it increases in the presence of *E*-2-hexenal. **(C)** *D. erecta* displays different turn rates and modulation of turn rates in the presence of ethyl acetate and *E*-2-hexenal. Differences of overall turn rates of odors with unstimulated condition, as well as modulation of the turn rate up and down the gradient (*when- to-turn*). The differences between turn rates are quantified with a two-sample *t*-Test (*p≤0.05, **p≤0.01, ***p≤0.001, ns, not significant *p*>0.05). **(D)** High proportions of turns towards the odor gradients are achieved in the presence of both ethyl acetate and *E*-2-hexenal compared to no- odor control. Differences of turning proportions with chance were tested by a one-sample one- sided *t*-Test (\**p*≤0.05, \*\**p*≤0.01, \*\*\**p*≤0.001, ns, not significant *p*>0.05). Differences of turning proportions of odors with unstimulated condition were tested by a two-sample *t*-Test (*p≤0.05, **p≤0.01, ***p≤0.001, ns: not significant *p*>0.05). **(E)** Both ethyl acetate and *E*-2-hexenal elicit strong weathervaning responses in *D. erecta* compared to no-odor control. In the boxplot of panels 7C and 7D, the mean (white line) is surrounded by a gray area that corresponds to 1.96 SEM (95% confidence interval), and a colored interval that corresponds to 1 SD. In graph of panels 7B and 7E, the mean is surrounded by the SEM displayed as a shaded area.

The ability of a larva to turn toward the odor gradient (*where-to-turn-to*) is a fundamental sensorimotor routine underlying accurate chemotaxis (Figure 6Ai). When exposed to their HOP odor, each species displayed a higher proportion of turns oriented towards the odor source. This bias was not specific to the HOP odor: it was observed for virtually every tested odor that elicited significant chemotaxis (Figure 6Aiii). Thus, the *where-to-turn-to* routine is not odor-specific or species-specific: it is a sensorimotor routine conditioning strong chemotaxis in all species.

Finally, we examined the contribution of *weathervaning* to orient in odor gradients (Figure 6Bi). This navigational strategy is achieved through smooth reorientation resulting in the continuous correction of the animal’s heading to improve its alignment with the local odor gradient. In Figure 6Bii, *weathervaning* is quantified as a modulation of the instantaneous reorientation rate by the bearing with respect to the odor gradient. The S-shape of the function implies that larvae tended to steer their runs toward the left side when the odor gradient pointed to the left, and vice versa. We found that all species implemented *weathervaning* upon HOP stimulation (Figure 6Bii). As for the other reorientation routines, this result could be generalized to a majority of the tested odors (Figure 6Biii). Nonetheless, we noted that the strength of the *weathervaning* was species-dependent. *D. erecta* displayed strong *weathervaning* for most odors whereas the bias implemented by *D. biarmipes* tended to be weak.

Since a given odor did not produce the same level of attraction in each species (Figure 2E), we further examined chemotaxis elicited by the set of HOP odors, which produced the most attractive behaviors in each species. Therefore, the comparative analysis based on the HOP behavior provided information about the characteristic sensorimotor strategies implemented by each species for effective chemotaxis. In the rest of the analysis, we extended the HOP comparison to special case studies that enabled an assessment of the conservation of navigational strategies between attractive odors within one species (Figure 7), as well as differences in the responses elicited by the same odor across species (Figures 8 and 9).

**Figure 8:**
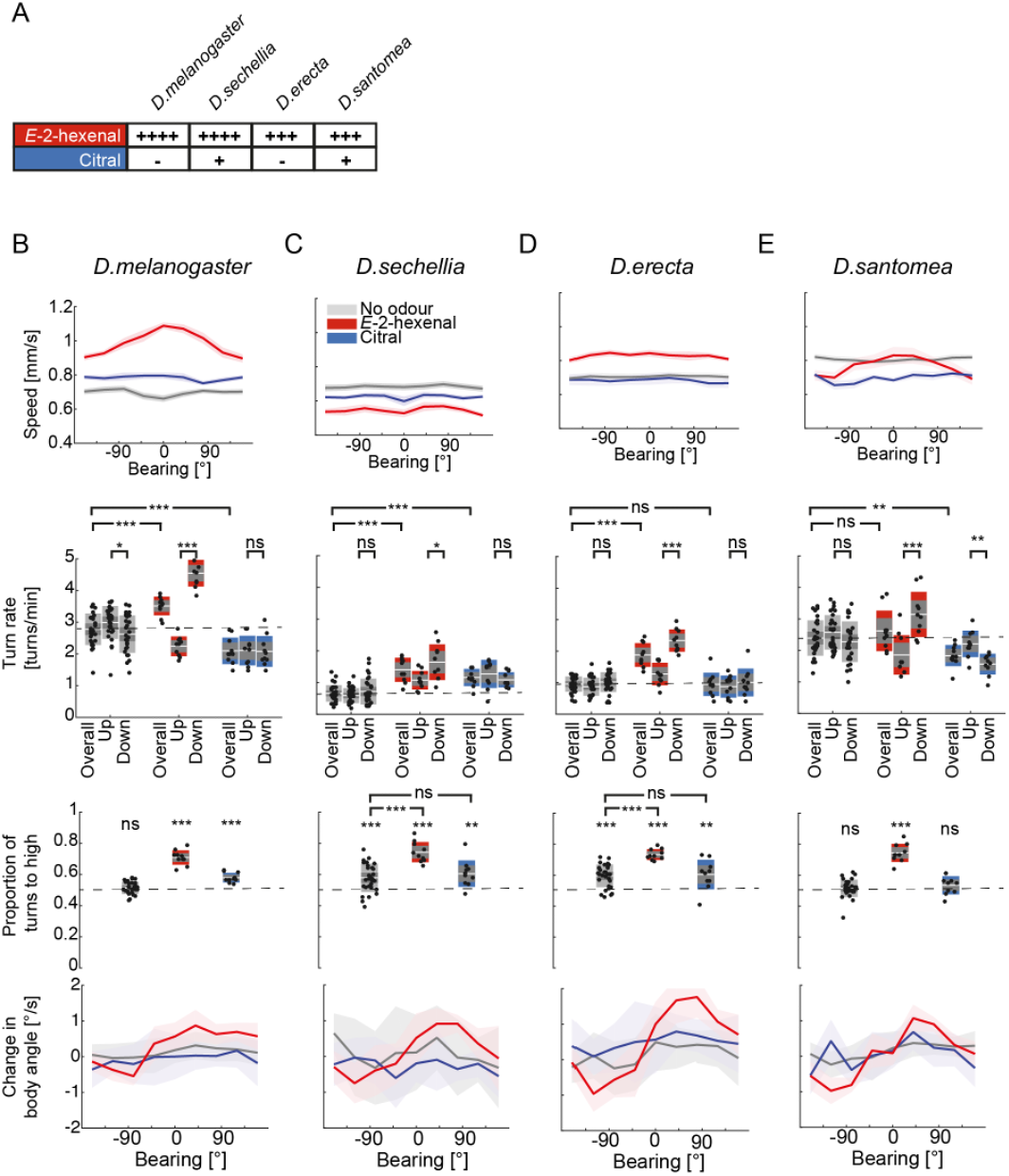
Similar attraction to the same pair of odors can result from different sensorimotor strategies across species. **(A)** *E*-2-hexenal and citral elicit strong (high PIs) and weak (low PIs) attractive responses, respectively, in *D. melanogaster*, *D. sechellia*, *D. erecta* and *D. santomea*. Strong attraction to *E*-2-hexenal and weak attraction to citral result from differences in the run speed and run speed modulation, the turn rate and turn rate modulation, the direction of turns and weathervaning in *D. melanogaster* **(B)**, *D. sechellia* **(C),** *D. erecta* **(D)** and *D. santomea* **(E)**. Differences of overall turn rates of odors with unstimulated condition, as well as differences of turn rates were tested by a two-sample *t*-Test (*p≤0.05, **p≤0.01, ***p≤0.001, ns: not significant *p*>0.05). Differences of turning proportions with chance were tested by a one-sample one-sided *t*-Test (\**p*≤0.05, \*\**p*≤0.01, \*\*\**p*≤0.001, ns: not significant *p*>0.05). Differences of turning proportions of odors with unstimulated condition were tested by a two-sample *t*-Test (*p≤0.05, **p≤0.01, ***p≤0.001, ns: not significant *p*>0.05). Number of trials per experimental condition (species — odor): 10. In the boxplots of the figure, the mean (white line) is surrounded by a gray area that corresponds to 1.96 SEM (95% confidence interval), and a colored interval that corresponds to 1 SD. In graph of the speed and reorientation rate, the mean is surrounded by the SEM displayed as a shaded area.

**Figure 9:**
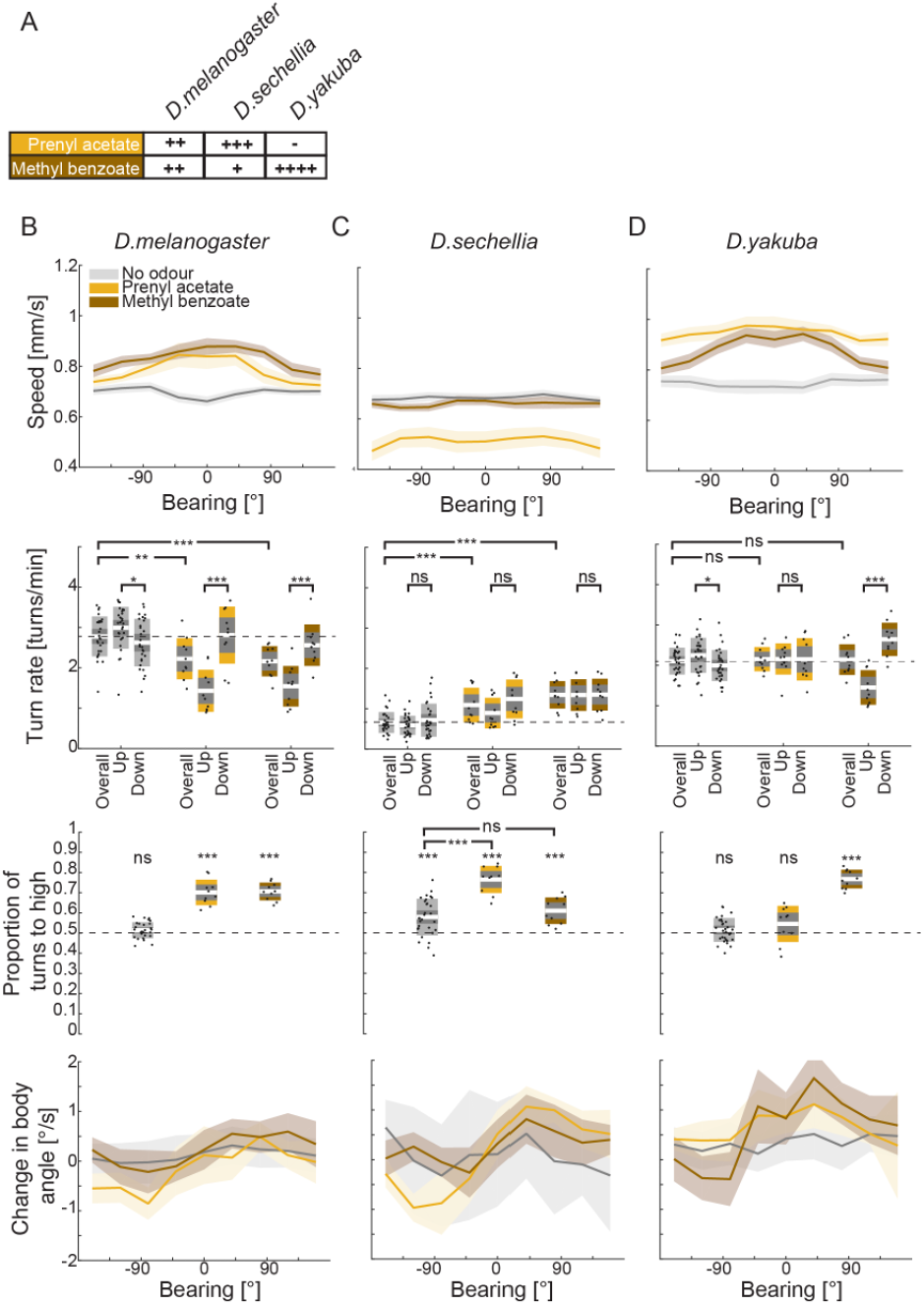
Inter-species variability: distinct attraction to the same pair of odors across species and are under different sensorimotor control. **(A)** Prenyl acetate and methyl benzoate elicit variable responses in *D. melanogaster*, *D. sechellia*, *D. yakuba*. **(B)** The same level of attraction to prenyl acetate and methyl benzoate elicit similarities in the modulation of run speed, turn rate, direction of turns and weathervaning in *D. melanogaster*. **(C)** Opposing behavioral trends, strong attraction to prenyl acetate and weak attraction to methyl benzoate elicit differences in the run speed and run speed modulation, the turn rate and turn rate modulation, the direction of turns and weathervaning in *D. sechellia*. **(D)** Opposing behavioral trends, weak attraction to prenyl acetate and strong attraction to methyl benzoate elicit differences in the modulation of the run speed and turn rate, the direction of turns and weathervaning in *D. yakuba*. Differences of overall turn rates of odors with unstimulated condition, as well as differences of turn rates up and down for each condition were tested by a two-sample *t*-Test (*p≤0.05, **p≤0.01, ***p≤0.001, ns: not significant *p*>0.05). Differences of turning proportions with chance were tested by a one-sample one-sided *t*-Test (\**p*≤0.05, \*\**p*≤0.01, \*\*\**p*≤0.001, ns, not significant *p*>0.05). Differences of turning proportions of odors with unstimulated condition were tested by a two-sample *t*-Test (*p≤0.05, **p≤0.01, ***p≤0.001, ns: not significant *p*>0.05). Number of trials per experimental condition (species — odor): 10. In the boxplots of the figure, the mean (white line) is surrounded by a gray area that corresponds to 1.96 SEM (95% confidence interval), and a colored interval that corresponds to 1 SD. In the graph of the speed and reorientation rate, the mean is surrounded by the SEM displayed as a shaded area.

While the HOP odor of each species elicited a unique modulation of the sensorimotor routines, this modulation varied within a species. Different odors could elicit different parametric arrangements of the sensorimotor routines, resulting in the same overall attraction level. This point is illustrated in Figure 7A by comparing the olfactory responses of *D. erecta*. While *D. erecta* larvae were strongly attracted to both ethyl acetate and *E*-2-hexenal (a high preference index marked as “+++”), the baseline run speed was higher for *E*-2-hexenal compared to ethyl acetate (Figure 7B). Consistent with the results of the HOP odor (2- heptanone) of *D. erecta*, ethyl acetate and *E*-2-hexenal did not elicit orientation-dependent run speed modulation (Figure 7B). While ethyl acetate did not elicit orientation-dependent turn rate modulation (*when-to-turn*), *E*-2-hexenal did (Figure 7C). Finally, both odors induced a higher proportion of turns towards the odor (*where-to-turn-to*) and marked *weathervaning* (Figure 7D-E).

Based on the results of Figure 7, we conclude that the effective localization of an odor source does not critically depend on the run speed modulation and the *when-to-turn* routine. The biases in the direction of turning and *weathervaning* appear to be sufficient to ensure strong chemotaxis in *D. erecta*. It is possible that the increase in run speed and/or the enhanced *weathervaning* elicited by ethyl acetate compensate for a loss in the *when-to-turn* modulation. This behavioral comparison between two strongly attractive odors suggests that losing the *when-to-turn* routine (control of timing of stop-turn) does not necessarily abolish chemotaxis.

In Figures 5 and 6, we established that each HOP odor elicited a species-specific sensorimotor strategy that modulated the baseline run speed and turn rate. Despite these differences, three sensorimotor routines that control turning — *when-to-turn*, *where-to-turn-to* and *weathervaning* — were universally implemented by all tested species when stimulated by the HOP odors. In Figure 8, we asked whether this conclusion held true for other odors that elicited strong chemotaxis in a subset of species. The strong attraction of *D. melanogaster*, *D. sechellia*, *D. erecta* and *D. santomea* to *E*-2-hexenal (Figure 8A) was underpinned by an equal modulation of the sensorimotor routines *when-to-turn*, *where-to-turn-to*, and *weathervaning* to chemotax toward the odor (Figure 8B-E). Conversely, a modulation of the run speed as a function of the animal’s orientation was clearly implemented by *D. melanogaster* and *D. santomea*, whereas it was weak or inexistent for *D. sechellia* and *D. erecta* (Figure 8, top row).

In *D. melanogaster*, *D. sechellia*, *D. erecta* and *D. santomea*, citral elicited weak or no attraction (Figure 8A). Compared to *E*-2-hexenal, citral produced a very weak or insignificant turning bias (*where-to-turn*) towards the gradient (Figure 8, third row). A low turning bias did not appear to be sufficient to direct reliable chemotaxis in *D. melanogaster* (Figure 8B, third row). Similarly, citral produced virtually no *weathervaning* to locate the odor source across species (Figure 8, fourth row). The run speed modulation displayed by *D. melanogaster*, *D. erecta* and *D. santomea* for the attractive odor *E*-2-hexenal was not implemented in response to citral, even though citral produced a baseline run speed that is unique to each species.

Finally, citral did not elicit orientation through the *when-to-turn* routine in any of the species (Figure 8, second row), except for *D. santomea* where an unusual inversion produced more turning during up-gradient than down-gradient movements.

Figure 8 yields two separate conclusions. First, strong attraction to the same odor, *E*-2- hexenal, can result from a similar strategy across species: a strong sensory modulation of the *when-to-turn*, *where-to-turn-to* and *weathervaning* routines. Second, the weak recruitment of one sensorimotor routine does not guarantee the emergence of reliable chemotaxis. This conclusion is drawn from the example of citral, which produced a change in run speed and a turning bias without promoting attraction to the odor source.

While many odors elicited attraction levels that were relatively conserved across species (e.g., *E*-2-hexenal, Figure 8), some odors elicited heterogeneous behavioral responses. This inter- species variability is illustrated in Figure 9 by the odor pair prenyl acetate and methyl benzoate. We found a remarkable qualitative change in the attraction level elicited by prenyl acetate in *D. sechellia* (strong attraction) and *D. yakuba* (virtually no response) (Figure 9A). By contrast, methyl benzoate was strongly attractive in *D. yakuba*, but weakly attractive in *D. sechellia*. The opposed preferences for prenyl acetate and methyl benzoate did not, however, indicate a switch in the underlying sensorimotor controls.

In response to prenyl acetate, *D. sechellia* modulated neither the run speed nor the *when-to-turn* sensorimotor routine to achieve attraction (Figure 9C, top two rows). Instead, strong attraction to prenyl acetate emerged from a marked use of the *where-to-turn-to* and *weathervaning* routines. The weak attraction to methyl benzoate resulted from a loss of the *where-to-turn-to* sensorimotor control and a strong reduction in *weathervaning*. Meanwhile attraction to methyl benzoate in *D. melanogaster* and *D. yakuba* correlated with a modulation of the run speed, the *when-to-turn* and the *where-to-turn-to* sensorimotor routines (Figure 9B and 9D). The absence of attraction to prenyl acetate found in *D. yakuba* correlated with a loss of the run speed modulation, *when-to-turn*, *where-to-turn-to*, and *weathervaning* routines. Unlike *D. sechellia* and *D. yakuba*, attractive behavior to prenyl acetate and methyl benzoate did not involve strong *weathervaning* in *D. melanogaster* (Figure 9B, bottom row). This result was consistent with the fact that *weathervaning* was less pronounced in *D. melanogaster* than in most other species (Figure 6B).

All in all, we conclude from Figure 9 that the *where-to-turn-to* control is critical to ensure strong chemotaxis as shown in the behavioral switch of *D. sechellia* and *D. yakuba* in response to prenyl acetate and methyl benzoate. Run speed modulation and the *when-to-turn* control are not strictly necessary to produce strong chemotaxis as seen in the case of prenyl acetate in *D. sechellia*, where the absence of stopping bias while moving down-gradient can be compensated by strong *where-to-turn-to* control paired with *weathervaning*.

## Discussion

How have neural functions evolved to accommodate specific behavioral needs associated with the search for food? To answer this question, one can focus on exploring the structural and functional relationships of the brain in a particular model organism. While this approach has advanced our understanding of the design principles of neural circuits underpinning vision in fruit flies and mice (Borst and Helmstaedter 2015), it has also been hampered by two obstacles: the numerical complexity of even the “simplest” brains, and major limitations in our understanding of the behavioral repertoire animals have evolved to thrive in an ecosystem. Moreover, the study of olfactory functions in one model organism, adapted to a particular ecological niche, is unlikely to explain adaption of food-search behavior in a vastly different setting such as in the lushness of a tropical forest or the harsh scarcity of a desert.

A complementary approach to unravel brain functions is to exploit the behavioral variability observed in closely related species to disentangle the link between genetic and structural changes in neural circuits and their specialized functions. This approach has clarified the genetic basis of variability in burrowing behavior and parental care in wild mice (Weber, Peterson et al. 2013). Although neurobiology has been dominated by studies focused on the detailed inspection of a handful of model organisms in recent decades, the advent of new techniques such as CRISPR/Cas9 to characterize and modify the genomes of non-traditional model systems has opened unprecedented opportunities for comparative analysis in systems neuroscience (Matthews and Vosshall 2020).

Invertebrates represent a system of choice to study the genetic and neural bases of behavioral adaptation due to the (more) tractable number of neurons of their nervous systems. To adapt to different ecological niches and environmental conditions, the chemosensory systems of insects have undergone changes on a more rapid timescale than other gene families (Sánchez-Gracia, Vieira et al. 2009). Accordingly, olfaction has been a productive testbed for identifying the genetic and neural substrates of evolutionary specialization in insects (Hansson and Stensmyr 2011, Zhao and McBride 2020). In the present work, we conducted a systematic comparison of olfactory behaviors in the genus Drosophila. We focused on the sensorimotor routines directing chemotaxis at the larval stage (Figure 1). In *D. melanogaster*, chemotaxis is controlled by a process of active sensing in which changes in odor concentration are detected through movements of the olfactory organs during runs and lateral head sweeps (head casts) (Gomez- Marin and Louis 2012). Here, we characterized how the organization of the sensorimotor routines — the algorithm — underlying larval chemotaxis has evolved in eight Drosophila species. This algorithmic comparison is a first step toward a mechanistic inspection of how behavioral specialization has emerged from changes in neural circuits.

### Behavioral evolution arises from changes in peripheral and central circuits

Two complementary mechanisms have been proposed to explain the emergence of specialized olfactory behaviors: changes in the peripheral sensory system and changes in higher-order circuits of the central brain (Zhao and McBride 2020). According to the first mechanism, the receptive field of a species can be dramatically altered by changes in the expression level and the coding sequences of odorant receptor (OR) and ionotropic receptor (IR) genes. Olfactory sensitivity can also be modified by increasing the number of olfactory sensory neurons (OSNs) expressing a particular OR. Such changes can modify the saliency of olfactory objects associated with food sources or danger. Evolutionary changes in the peripheral tuning of the olfactory system have been uncovered in *D. sechellia*, which feeds and breeds specifically on the noni fruit — a substrate that is toxic to all other Drosophila species. This specialization to a unique ecological niche results from an increase in the number of OSNs that express the Or22a receptor tuned to esters emitted by the noni fruit (Dekker, Ibba et al. 2006, Auer, Khallaf et al. 2020). In addition, the Or22a receptor of *D. sechellia* exhibits a heightened sensitivity to the noni fruit compared to its sibling species, *D. melanogaster* and *D. simulans*. Moreover, changes at the level of the first-order olfactory sensory neurons are accompanied by a modification of the structure of second-order project neurons. In the seasonal specialist *D. erecta*, species-specific host-seeking behavior has resulted from the overrepresentation of a dedicated class of OSNs tuned to the pandanus tree (Linz, Baschwitz et al. 2013).

Behavioral specialization has also been linked to changes in central olfactory circuits in the context of mating. During courtship, flies ignore mating partners from heterospecific species. Targeting a suitable mate entails the recognition of sex pheromones in addition to species-specific steps in love parades composed of elementary forms of social interactions (Sokolowski 2001). Notably, the pheromone 7,11-heptacosadiene promotes courtship in *D. melanogaster* whereas it suppresses this behavior in the sibling species *D. simulans*. The differential response to the pheromone does not result from a specialization of the peripheral olfactory system, but a change in the balance between excitatory and inhibitory inputs that control a central circuit promoting courtship (Seeholzer, Seppo et al. 2018). In a separate study, mate discrimination in *D. mojavensis* has been shown to arise from the differential interpretation of the same peripheral cues between different subspecies (Khallaf, Auer et al. 2020). In a similar way, the species-specificity of courtship follows from the response to selective features in the organization of courtship songs through which males attract females of the same species. In a genetic screen, the ability of *D. simulans* and *D. mauritiana* to differentiate each other has been linked to a calcium-activated potassium channel expressed in a variety of neurons of the nervous system (Ding, Berrocal et al. 2016). These findings led to the idea that evolution can adjust behavior by modifying the functional properties of hubs in neural circuits. Together, specialization in courtship behavior can arise from changes in the peripheral olfactory system, signal processing units and motor pathways (Sato, Tanaka et al. 2020). The relevance of each level of regulation remains to be determined.

By comparing the behavioral responses elicited by a common set of 19 odors (Figure 2E), we delineated the features of the olfactory navigation algorithm that are conserved and variable in eight species of the genus Drosophila. Our quantitative analysis shed light on two types of evolutionary changes: (i) the capacity to detect an odor (tuning of the peripheral olfactory system) and (ii) alterations in the sensorimotor routines underlying orientation maneuvers. While the first type of changes might result from transformations in the expression of OR genes or their coding sequences, the second type is likely to entail the modification of higher-order neural circuits. Using automated tracking of groups of freely moving larvae (Swierczek, Giles et al. 2011) and statistical correlations between behavior and sensation (Gomez-Marin, Stephens et al. 2011, Gershow, Berck et al. 2012), we carried out a thorough screen of the hierarchy of sensorimotor routines that underlies larval chemotaxis (Figure 1). At the onset of the screen, nothing allowed us to predict the degree of inter-species behavioral variability: it was plausible that the navigation algorithm characterized in *D. melanogaster* represented an optimal solution shared with all other Drosophila species. While the basic navigation strategy observed in *D. melanogaster* appears to be largely conserved within the genus Drosophila, significant variability was found both in the receptive fields (Figure 2E) and the organization of elementary sensorimotor routines into species-specific algorithms that promote strong chemotaxis (Figure 10A).

**Figure 10:**
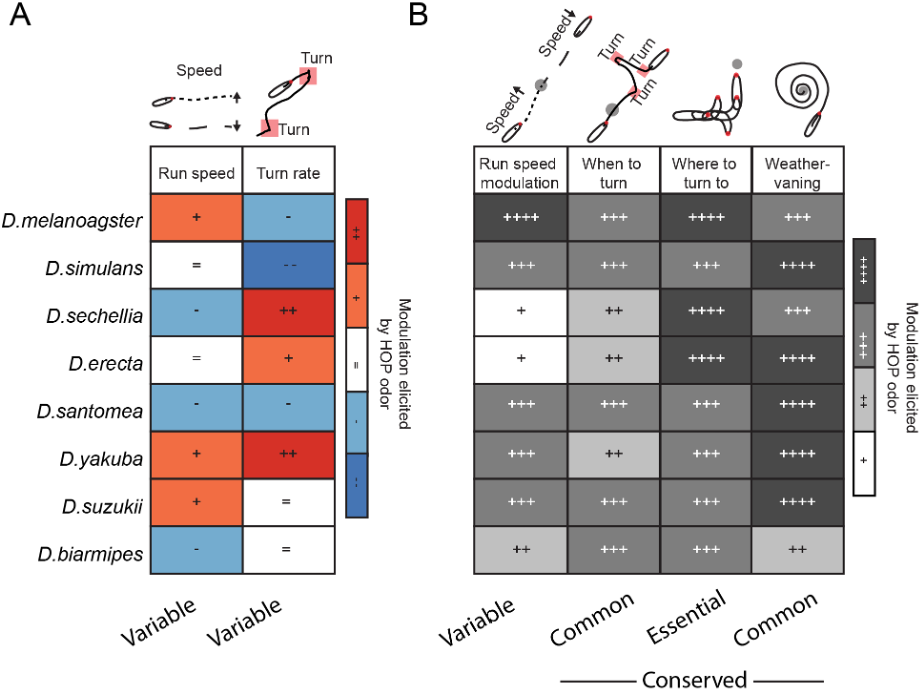
Summary table of the contributions of different sensorimotor routines to navigational strategies across species. Modulation strength is denoted by a -,+ or = as elicited by the high-olfactory-preference (HOP) odor of each species. The modulation levels are qualitatively color-coded based on the sign and magnitude of the change induced by the odor. The color-coding does not convey any information about the statistical significancy of differences between the no-odor and odor conditions.

### Responses to a common set of 19 odors differ across species of the genus Drosophila

When reduced to a preference index that measures the level of attraction to the source, the behavioral profile of each species —a column in the table shown in Figure 3E— suggests that each species responds to an idiosyncratic set of odors. An odor can produce attraction in one species while eliciting no significant response in its sister species. Consider for instance prenyl acetate in *D. santomea* (strong attraction) and *D. yakuba* (no response). The simplest explanation for this all-or-none behavioral difference is that *D. yakuba* might have undergone detrimental changes in one or multiple OR genes crucial for the detection of prenyl acetate. The same might explain why *D. sechellia* is virtually not responsive to methyl acetate whereas *D. melanogaster* and *D. simulans* are strongly attracted by this odor. Given that Or42a and Or42b display the highest sensitivity to methyl acetate (Münch and Galizia 2016), one can speculate that evolutionary changes specific to these two OR genes might be responsible for the loss of attraction of *D. sechellia* to methyl acetate. The same two ORs might be implicated in the heterogeneous responses elicited by methyl benzoate (Figure 3A). More generally, genetic changes in a subset of OR genes are expected to, at least partly, explain the inter-species behavioral variability observed in the group of “medially attractive” odors defined in Figure 3B (yellow color). By contrast, some odors such as ethyl acetate and 1-hexanol elicit attraction in all species (Figure 2E and Supplementary Figure 1). Other odors elicit no response in nearly all species (Figure 3B, blue color) or, in some cases, even active repulsion reflected in a faster increase of the distance to the source compared to the no-odor condition (e.g., acetic and octanoic acids in *D. erecta*, Supplementary Figure 2). We hypothesize that odorant molecules eliciting a universally strong response might fulfil a common role in the ecological niches of larvae of the genus Drosophila, which must have led to the conservation of the OR repertoire that permits their detection.

The inter-species comparison of olfactory behaviors reveals a remarkable correlation between the molecular weight of an odorant molecule and the consistency of its level of attraction across species. Molecules with a low molecular weight tend to produce strong attraction and limited variability across species. Molecules with medial molecular weights (120-140 g/mol) yield higher variability in their levels of attraction (Figure 3C). Molecules with high molecular weights (>140 g/mol) produce weak or no significant attraction — acetic acid representing the notable exception to this rule: despite its small size, it does not produce strong attraction. This exception of acetic acid is likely to be explained by the toxicity of acetic acid (Devineni, Sun et al. 2019).

To our knowledge, there is no explanation for the observed correlation between the molecular weight of an odor and the degree of inter-species variability this odor elicits behaviorally. While it is plausible that the reduced diffusion of heavier odorant molecules could create larger fluctuations in the profile of their odor gradients, this factor is unlikely to produce significant behavioral variability across species since the concentration profiles of gradients created with molecules having different molecular weights are very similar (Louis, Huber et al. 2008).

Are the response profiles of pairs of closely related species (e.g., *D. melanogaster* and *D. simulans*), more similar than those of more remote species (e.g., *D. melanogaster* and *D. suzukii*)? We tested this hypothesis by building a dendrogram — a behavioral tree — based on the Euclidean distance between pairs of response profiles. While the behavioral tree (Figure 3G) reproduces important aspects of the commonly-accepted phylogenetic tree (Figure 3F), the position of *D. sechellia* and *D. yakuba* in the tree stands out as exceptions. This result is not surprising for *D. sechellia* given that it has evolved as an extreme specialist. By contrast, one might not have anticipated strong behavioral divergences in olfactory behaviors between *D. santomea* and *D. yakuba*. In addition, it is notable that *D. biarmipes* —a species whose behavior remains poorly characterized— stands out as a clear outlier in the behavioral dendrogram. It is unknown why *D. biarmipes* would have lost its ability to efficiently chemotax to most odors tested in the screen.

### Species-specific variability in locomotor speed and the control of turning maneuvers

As illustrated in Figure 1, two main sensorimotor variables underlie the control of larval chemotaxis: (i) run speed quantified through the speed of the larval midpoint, and (ii) active sampling quantified through the amplitude of lateral head movements and turn rate (Figure 4A). Although larvae of the eight species considered in the present study (Figure 2A) do not show major morphological differences, their locomotor behaviors vary considerably, particularly with respect to their turn rates (Figure 4B). This observation is consistent with the results of a recent screen of larval locomotion in the genus Drosophila (Matsuo, Nose et al. 2020). In unstimulated conditions, species can be grouped into “non-turners” and “turners” (Figure 4B, left). This grouping is conserved when larvae are stimulated by their high-olfactory-performance (HOP) odor (Figure 4B, right). The reliability of the classification was established by a linear discriminant analysis. The odor stimulation converts a slight correlation between run speed and the turn rate into a slight anticorrelation between these variables. The “non-turner” group includes *D. sechellia*, *D. erecta* and *D. biarmipes*. Notably, *D. sechellia* and *D. erecta* are part of two different complexes (*D. sechellia* belongs to the simulans complex whereas *D. erecta* belongs to the yakuba species complex). We observe a similar split in the two Asian species since *D. suzukii* is a turner while *D. biarmipes* is a non-turner. Therefore, being a turner or a non-turner is not a behavioral property that is shared among species that are closely-related phylogenetically.

Next, we explored in more detail the modulatory effects of HOP stimulation on the run speed and the turn rate independently. Stimulation by the HOP odor produced either an increase, a decrease or no change in run speed compared to the no-odor stimulation condition (Figure 4C). As outlined in Figure 10A, three species demonstrated an acceleration: *D. melanogaster*, *D. yakuba* and *D. suzukii*. These species are part of different phylogenetic complexes. Likewise, a deceleration is observed in three species that are separate phylogenetically: *D. sechellia*, *D. santomea* and *D. biarmipes*. Stimulation by the HOP odor produced no change in the average run speed of *D. simulans* and *D. erecta*. These results illustrate a split of the simulans, the yakuba and the suzukii complexes.

No obvious correlation is found between the changes in run speed and turn rate resulting from the exposure to the HOP odors (Figure 10A). Most species demonstrate either an increase or no change in the overall turn rate. The only species characterized by a strong decrease in turn rate is *D. simulans*. Each species of the yakuba complex shows a correlation in the change in run speed and turn rate: both variables increase in *D. yakuba*, whereas they decrease in *D. santomea*. While *D. santomea* stands out as the species that is attracted to the largest number of odors and with the highest persistence (Figure 2E and Supplementary Figure 2), the higher accuracy of its olfactory behavior does not result from faster runs or more turning, but it shows the opposite trend. The complex relationship between run speed, turn rate and the strength of chemotaxis underscores the importance of comparing the elementary sensorimotor routines directing larval chemotaxis (Figure 1).

### Species-specific modulation of run speed and turn rate by the olfactory experience

Our quantitative analysis of larval chemotaxis indicates that run speed can be instantaneously modulated by the animal’s orientation with respect to the gradient (animal’s bearing). Typically, an acceleration is observed for upgradient runs while a deceleration follows downgradient runs. This phenomenon has been reported for *D. melanogaster* in earlier work (Gershow, Berck et al. 2012, Schleyer, Reid et al. 2015). The modulation of the run speed by the bearing is found in 5 out of the 8 species: it is most pronounced in *D. melanogaster* and *D. yakuba*. It is not universal though (Figure 10B): in *D. sechellia*, for instance, an overall decrease in run speed takes place without any selective modulation as a function of the bearing (Figure 5Aii). The absence of modulation is not specific to the stimulation of *D. sechellia* by its HOP odor: it applies to most of the 19 tested odors (Figure 5Aiii). In future work, it will be interesting to define how the bearing to the gradient is detected during runs and how this variable modulates the run speed.

Whichever mechanism underlies this sensorimotor routine, it highlights that changes in odor concentration do more than simply control the probability of switching between running and stopping/turning: the detection of concentration derivatives can also continuously adjust the speed of locomotion (Figure 1Bi).

Turn rate is another locomotor response that is strongly modulated by the animal’s bearing in an odor gradient. Larvae tend to turn more during movement away than toward the gradient. This modulation forms the basis of the *when-to-turn* sensorimotor control (Figure 5Bii). Remarkably, this sensorimotor routine appears to be independent of the net effect that the odor has on the overall turn rate. For some species, such as *D. sechellia*, HOP stimulation produces an increase in turn rate, whereas it yields either no change or an overall decrease in turn rate in other species such as *D. simulans*. Even the species classified as non-turners (Figure 4B) modulate their turn rate as a function of the bearing. In contrast with the species-specific modulation of the run speed, the *when-to-turn* modulation of the turn rate appears to be universal (Figure 10B).

The conservation of this sensorimotor routine across species of the genus Drosophila is consistent with its use in more primitive organisms like bacteria where chemotaxis is achieved by suppressing turning while moving upgradient of food-related substances (Sourjik and Wingreen 2012).

What controls the universal modulation of the turn rate as a function of the bearing while allowing for species-specific differences in the effects of the odor induced modulation of the overall turn rate? In particular, what explains that the overall turn rate increases in response to ethyl butyrate in *D. melanogaster* and *D. sechellia* while it decreases in *D. simulans* (Supplementary Figure 4B)? Even though the neural substrates of most sensorimotor routines remain unknown in the *Drosophila* larva, a critical aspect of the control of turn rate has been mapped in *D. melanogaster* onto a descending neuron, PDM-DN, which promotes stopping in response to negative gradients measured mainly by two olfactory sensory neurons expressing the *Or42a* and *Or42b* genes (Tastekin, Khandelwal et al. 2018). Ethyl butyrate activates both *Or42a* and *Or42b* OSNs. By a mechanism that is left to be elucidated, circuits that process the peripheral activity of the *Or42a* and *Or42b* OSNs upstream from PDM-DN must have evolved to produce an overall downregulation in turn rate in *D. simulans* compared to an overall upregulation in *D. melanogaster* and *D. sechellia* (Figure 10A and Supplementary Figure 4B).

Despite this change, each of these three species is capable of robust *when-to-turn* modulation by promoting turning downgradient while suppressing turning upgradient (Figure 5Biii). Whether and how changes in the PDM-DN circuit contributes to these evolutionary differences within the simulans complex is an intriguing question left to be addressed in future work.

### The ability to turn toward the gradient is shared by all species of the genus Drosophila

Through the *when-to-turn* sensorimotor routine, all species can adjust their turn rates based on their sensory experience (Figure 5). The strength of the sensory modulation is nonetheless variable across species and odors. By contrast, all species are characterized by a robust *where-to-turn-to* routine (Figure 1Biii), reflected in a strong bias of their turns toward the gradient of their HOP odors (Figure 6Bii). Moreover, the *where-to-turn-to* routine shows little variability across odors (Figure 6Aiii). Larvae from all species perform smooth continuous turns — *weathervaning* (Figure 1Biv) — toward the HOP gradient (Figure 6Bii), but with a higher degree of variability across species and odors (Figure 6Biii). In conclusion, *when-to-turn*, *where-to-turn- to* and *weathervaning* represent sensorimotor routines shared by all tested species to achieve chemotaxis. In spite of this similarity, more cases of species- and odor-specific differences are observed for the *when-to-turn* and *weathervaning* routines compared to the *where-to-turn-to* routine. Intriguingly, *weathervaning* is not a sensorimotor routine that is strongly expressed in *D. melanogaster* where it was first described (Gomez-Marin and Louis 2012).

A striking correlation is found between the inability of larvae to chemotax toward specific odors, butyl butyrate, citral and octanoic acid (Figure 2E) and the absence of turning bias — *where-to-turn-to* modulation — for these odors. Together with the inspection of variability in chemotactic responses to specific odors (Figures 7-9), we conclude that the *where-to-turn-to* routine is essential to ensure that the navigation algorithm produces strong odor attraction. Species-specific losses in the *where-to-turn-to* routine for a subset of odors also reveals a relationship with the *when-to-turn* routine. Consider the significant attraction that 2-heptanone elicits in *D. santomea* but not *D. yakuba* (Figure 2E). This loss of attraction to 2-heptanone is correlated with the inability of *D. yakuba* to perform robust *when-to-turn* (Figure 5Biii) and *where-to-turn-to* modulation in response to this odor (Figure 6Aiii). We speculate that peripheral changes in the olfactory system of *D. yakuba* might prevent the detection of changes in the concentration of 2- heptanone. Similarly, a loss of attraction to methyl acetate in *D. sechellia* (Figure 2E) is correlated with the inability of this species to carry out any of the four basic sensorimotor routines (Figure 1B). Unlike 2-heptanone in *D. yakuba*, *D. sechellia* must be able to detect methyl acetate since this odor induces more turning overall (Supplementary Figure 4B) as well as a reduced distance to the source compared to the no-odor control (Supplementary Figure 2). Despite an increase in turning elicited by methyl acetate, *D. sechellia* is unable to turn toward this odor, which indicates either an inability to detect changes in the concentration of methyl acetate at the sensory periphery or a modification of central circuits leading to a change in the valence of this odor in *D. sechellia* specifically.

### Different ways to achieve robust chemotaxis within and across species

After having observed inter-species variability in the organization of the sensorimotor algorithm underlying chemotaxis (Figure 10), we evaluated the existence of algorithmic variability in the behaviors elicited by different odors within the same species. Two possible outcomes were envisioned. First, chemotaxis could result from a fixed combination of elementary sensorimotor routines that is specific to one species. According to this view, orientation responses could be expressed with different strengths depending on the nature and intensity of the odor, but the combination of sensorimotor routines would not vary. Second, odors that are processed differently at the sensory periphery could recruit distinct sensorimotor routines. We inspected the validity of the two alternatives in *D. erecta* by comparing the responses induced by two equally attractive odors: ethyl acetate and *E*-2-hexenal (Figure 2E and Figure 7A). We found that these two odors elicit different sets of sensorimotor controls (Figure 7B). Ethyl acetate does not elicit a strong *when-to-turn* modulation (prevalence of turning during downgradient) whereas *E*-2-hexenal does (Figure 7C). Thus, the *when-to-turn* routine is not necessary to ensure chemotaxis. On the other hand, *E*-2-hexenal induces an increase in the overall run speed that is not observed for ethyl acetate (Figure 7B). The *where-to-turn-to* and *weathervaning* routines are common to the response elicited by both odors, underscoring the importance of biasing turns toward the gradient for flies to perform strong chemotaxis (Demir, Kadakia et al. 2020).

The results of the odor comparison of Figure 7 indicate that there is more than one way to achieve strong chemotaxis. We speculate that two odors might direct qualitatively different navigation algorithms by recruiting the activity of distinct descending pathways, which would differentially tune the basic sensorimotor routines underlying larval chemotaxis (Figure 1B). This hypothesis is supported by the observation that the two odors featured in Figure 7, ethyl acetate and *E*-2-hexenal, do not activate the same subset of larval OSNs in *D. melanogaster*. If we assume that the overall response profile of the larval OR repertoire is conserved between *D. melanogaster* and *D. erecta*, ethyl acetate is expected to predominantly activate Or42a and Or42b (Kreher, Mathew et al. 2008). *E*-2-hexenal would mainly activate a different subset of ORs: Or7a, Or42a and Or67b (Münch and Galizia 2016). Therefore, it is possible that activity mediated by Or7a and/or Or67b direct the *when-to-turn* sensorimotor routine specifically in response to *E*-2-hexenal. These considerations suggest that different odors might recruit distinct subsets of OSN channels and their downstream descending pathways, which would result in orientation responses — navigation algorithms — that are odor-specific (Jung, Hueston et al. 2015, Newquist, Novenschi et al. 2016).

Given that qualitative differences in chemotactic algorithms were observed between *E*-2- hexenal and ethyl acetate in *D. erecta* (Figure 7), we asked whether the same type of variability is observed in response to a given odor, *E*-2-hexenal, across different species. We tackled this question in the three species that are most closely related to *D. erecta*: *D. melanogaster*, *D. sechellia* and *D. santomea* (Figure 8). In all four species, *E*-2-hexenal produces strong attraction (Figure 8A). Not surprisingly, each species shows a strong turning bias toward the gradient that results in *where-to-turn-to* and *weathervaning*. A bias of stop-turn during downgradient motion (*when-to-turn* modulation) is also observed in all four species. Contrary to the evolutionary conservation of the *when-to-turn* and *where-to-turn-to* modulation, the modulation of the run speed is variable: in response to *E*-2-hexenal, *D. melanogaster* and *D. erecta* accelerate whereas *D. sechellia* and *D. santomea* decelerate. These observations are consistent with the run speed modulation (Figure 1Bi) reported for the HOP odors in Figure 10.

### From attraction to aversion in response to the same odor in different species

After having examined how the same odor can produce strong attraction through different arrangements of the sensorimotor routines across species (Figure 8), we considered the opposite phenomenon in Figure 9: prenyl acetate elicits strong attraction in *D. melanogaster* and *D. sechellia*, whereas it elicits no attraction in *D. yakuba*. In fact, prenyl acetate is one of the few odors that produce active aversion, as judged from the larger distance to the source that larvae maintained compared to the no-odor control (Supplementary Figure 2). The sensorimotor analysis of Figure 9 establishes that the loss of attraction of *D. yakuba* to prenyl acetate correlates with the complete absence of any sensorimotor routine contributing to positive chemotaxis: there is no modulation in run speed with respect to the bearing, no *when-to-turn* nor *where-to-turn-to* modulation and nearly no *weathervaning*. And yet, the fact that prenyl acetate produces an overall increase in run speed indicates that this odor is perceived by *D. yakuba*.

While it is unclear whether the inability of *D. yakuba* to chemotax toward prenyl acetate results from peripheral changes in the expression or activity of a subset of OR genes, such differences would not produce a general impairment in chemotaxis since *D. yakuba* larvae are capable of strong attraction to methyl benzoate compared to *D. melanogaster* and *D. sechellia* (Figure 9). The enhanced attraction of *D. yakuba* to methyl benzoate correlates with the combination of strong *when-to-turn*, *where-to-turn-to* and *weathervaning* modulations. In addition, it features a marked modulation of the run speed as a function of the bearing (Figure 9D). Intriguingly, the strong attraction to methyl benzoate appears to be specific to the yakuba complex: it is shared between *D. yakuba* and *D. santomea* (Figure 2E). The ecological significance of the loss of attraction of *D. yakuba* to prenyl acetate remains to be elucidated.

### In search of evolutionary principles directing the adaptation of navigation behavior

Considerable progress has been made toward understanding the molecular and cellular bases of olfactory behaviors in *D. melanogaster* (Cobb 1999, Fishilevich, Domingos et al. 2005, Kreher, Mathew et al. 2008, Louis, Huber et al. 2008, Mathew, Martelli et al. 2013). Based on published work, a conceptual foundation of the sensorimotor computation directing larval chemotaxis has emerged in *D. melanogaster* specifically (Gomez-Marin, Stephens et al. 2011, Gershow, Berck et al. 2012, Gomez-Marin and Louis 2012, Hernandez-Nunez, Belina et al. 2015, Schulze, Gomez-Marin et al. 2015, Louis 2020). The question that initiated the present study is how relevant the combination of sensorimotor routines — the algorithm — directing larval chemotaxis in *D. melanogaster* is to all species of the genus Drosophila. Our correlative analysis of the sensory inputs and the behavioral responses observed for 19 odors and 8 species allow us to draw two main conclusions. First, the algorithmic framework proposed for *D. melanogaster* is generalizable to all species tested in this study (Figure 1). The basic sensorimotor routines directing larval chemotaxis are shared behavioral building blocks.

Second, the exact arrangement of the sensorimotor routines varies across odors and species. Although all species of the Drosophila genus appear to be chemotaxing in a way that is qualitatively similar, the algorithm implemented by *D. melanogaster* is not a “one size fits all” solution adopted by all its closely related species. At the most basic level, different species display a different basal turn rate and locomotor speed (Figure 4C). Moreover, substantial differences are observed in the way that a given species achieves chemotaxis in response to distinct odors (Figure 7). The degree of algorithmic variability observed across strains of the same species remains to be determined in future work.

Not all the basic sensorimotor routines available to the larva (Figure 1B) are equivalent in their contribution to generate strong attractive responses. Two sensorimotor routines show high inter- species variability: the modulation of the overall run speed and turn rate by the odor (Figure 10A). This modulation can be positive or negative without precluding strong chemotaxis. In addition, there is no obvious correlation between the effects of an odor on the overall run speed and turn rate. By contrast, the ability to modulate the instantaneous run speed, turn rate and turn direction as a function of animal’s bearing is largely conserved across the eight species tested here (Figure 10B). One routine in particular stands out as being critical to ensure robust chemotaxis: the ability to direct turns toward the odor gradient (*where-to-turn-to* routine, Figure 1Biii). In absence of turn bias toward the gradient, chemotaxis is generally impaired (see for instance Figure 9B-D). Interestingly, the ability of larvae and adult flies to turn toward the odor gradient emerges as essential to achieve strong chemotaxis in computational simulations (Davies, Louis et al. 2015, Demir, Kadakia et al. 2020). By contrast, the case study of Figure 7 illustrates that the ability to modulate the run speed (Figure 1Bi) and the turn rate (*when-to-turn*, Figure 1Bii) as function of the bearing is not necessary to ensure strong chemotaxis.

For a larva to turn toward the gradient (*where-to-turn-to* routine, Figure 1Biii), it must be capable of sensing minute odor gradients through the spatial displacements of its olfactory organs during rapid head casts lasting a characteristic time of 200 ms. Thus, the detection of concentration derivatives is essential to enable chemotaxis. One explanation for the inability of a species to turn toward the gradient of a given odor might be explained by a loss in sensitivity to temporal changes in the odor concentration, while still being able to perceive the presence of this odor itself. This hypothesis could explain why *D. yakuba* shows no bias in turning toward prenyl acetate (Figure 9D), even though its speed increases upon stimulation by this odor (Figure 9D and Supplementary Figure 4). This phenomenon is unlikely to be explained by the simple loss of an OR gene: it might involve changes in the physiology of the olfactory sensory neurons involved in the detection of concentration derivatives. It could also result from structural changes in the organization of the peripheral olfactory system, as has been observed in adult flies (Auer, Khallaf et al. 2020). On the other hand, we found drastic changes in odor behavior that are compatible with the loss of peripheral ORs responsible for the detection of specific odors. For instance, prenyl acetate and citral produce no attraction and no overall change of run speed in *D. erecta* (Figure 2E and Supplementary Figure 4A), which suggests a genuine insensitivity to specific odors that are detected by other species. In a similar way, the recruitment and loss of chemosensory genes is thought to explain the ability of larvae to recognize individuals of the same species (Mast, De Moraes et al. 2014, Del Pino, Jara et al. 2015).

Not all inter-species behavioral differences can be readily explained by the evolution of the peripheral encoding of odors. The effect of an odor on the overall run speed represents a notable example. While odor stimulation induces an acceleration of locomotion in some species, it produces a deceleration in other species (Figure 10A). The positive or negative effects of the odor on the locomotor speed is largely conserved within one species across the set of tested odors (Supplementary Figure 4A). This inter-species difference in behavior is likely to result from changes in the central circuitry in charge of tuning the locomotor activity of the larva.

Similar functional modifications might explain changes in the valence of an odor across species. Prenyl acetate provides a good illustration when compared between *D. sechellia* and D. *yakuba* (Figure 9). Another remarkable change in larval chemotaxis is the persistence with which larvae stay near an odor source. Each species tends to lose interest in an odor source that is not associated with food, but the timescale of this habituation is species-specific. As seen in Supplementary Figure 2, *D. santomea* stays near the source of most odors for more than 3 min. By comparison, *D. sechellia* and *D. erecta* lose interest in most odors much more quickly (< 100s). Such changes in olfactory habituation are expected to involve more than the recruitment or losses of OR genes.

Is the variability in larval chemotaxis representative of evolutionary changes that pertain to navigation to other sensory modalities? Have phototaxis and thermotaxis evolved in a similar way? How relevant is chemotaxis to other more complex odor-related behavior such as burrowing to find food (Godoy-Herrera 1986)? In the dig-and-dive assay (Kim, Alvarez et al. 2017), *D. melanogaster* and *D. suzukii* have been shown to differ dramatically in the organization of their odor-driven exploration, their response to hypoxia and the long-term habituation to the presence of an appetitive odor in the absence of food. Due to their critical importance for the survivability of animal, we speculate that aversive behavioral response triggered by potentially harmful substances might display reduced variability across species compared to food search that can be successful in a variety of ways. The present analysis should provide a solid quantitative starting point to explore the genetic basis that underlies changes in the characteristics of peripheral sensors and neural circuits involved in larval chemotaxis to adapt to changing environmental conditions. Such an analysis has already been initiated to study inter-species differences in desiccation behavior and cold resistance (Kellermann, Hoffmann et al. 2018). With its detailed algorithmic understanding in *D. melanogaster*, larval chemotaxis stands out as an excellent system to identify generic principles directing behavioral adaptation.

## Acknowledgement

We are deeply thankful to Biafra Ahanonu who conducted pilot experiments related to this project during a summer internship project in 2010. We are grateful to Alex Gomez-Marin, Nicolas Gompel, Morea Phillips, Benjamin Prud’homme and Nicole Voges for valuable inputs during the initial exploratory phase of this project, as well as the Louis and Simpson lab members for inputs on the manuscript. We thank Nicolas Gompel for generously providing many stocks used in this study. The project benefited from the financial and intellectual support from the EU FLiACT Initial Training Network. This work was supported by the ‘Centro de Excelencia Severo Ochoa 2013-2017’, the CERCA Programme/Generalitat de Catalunya, the EMBL/CRG Systems Biology Program and the University of California, Santa Barbara.

## Materials and methods

### Fly Stocks

Fly stocks were raised on standard cornmeal-agar molasses medium at 22°C, 60%–70% relative humidity, and kept in a 12 h dark-light cycle unless indicated otherwise. Wild-type flies were used for all 8 species tested in this study. *D. melanogaster* (Canton-S) was donated by Ilona Kadow, *D. simulans* was donated by Janelia, *D. sechellia* (Tucson TSC#14021-0248-25, BPNG#97), D. erecta (188.1 BPNG#212), *D. santomea* (São Tomé and Príncipe, STO.4 BPNG#96), *D. yakuba* (Liberia, Tucson TSC#14021-0261.01, BPNG# 95), *D. suzukii* (France) and *D. biarmipes* (Cambodia, Tucson TSC#14023-0361.01, BNPG#107) were all donated by Nicolas Gompel. *w*/*w*;+/+;*Orco*^2^/*Orco*^2^ transgenic anosmic flies were used for control experiments to determine appropriate odor concentrations.

### Behavioral assay and experiment

All behavioral experiments were performed with third instar foraging larvae, at room temperature. A light pad (Slimlite Lightbox, Kaiser) illuminated the arena from above creating uniform daylight conditions. The behavioral assay consisted of a 3% agarose (SeaKem LE Agarose, Lonza) surface coating (50 mL) the bottom part of a large Petri dish with a diameter of 15 cm (Sarstedt). Two transparent reinforcement rings, used as wells to load the odor, were fixed onto the inner surface of the Petri dish lid, at positions along the midline of the dish, equidistant to each other as well as to the edge of the lid. All 19 odors (methyl acetate CAS No. 79-20-9; ethyl acetate CAS No. 141-78-6; i-amyl acetate CAS No. 123-92-; prenyl acetate CAS No. 1191-16-; methyl butyrate CAS No. 623-42-7; ethyl butyrate CAS No. 105-54-4; butyl butyrate CAS No. 109-21-7; methyl hexanoate CAS No. 106-70-7; methyl benzoate CAS No. 93-58-3; Anisole CAS No. 100-66-3; g-octalactone CAS No. 104-50-7; 1-hexanol CAS No. 111- 27-3; 2-hexanol CAS No. 626-93-7; 2-heptanone CAS No. 110-43-0; E-2-hexenal CAS No. 6728-26-3; citral CAS No. 5392-40-5; acetic acid CAS No. 64-19-7; hexanoic acid CAS No. 142- 62-1; octanoic acid CAS No. 124-07-2) were purchased from Sigma-Aldrich and were presented to the larvae by loading two 10 µL droplets into the two transparent reinforcement rings fixed to the lid. Dilutions for all odors were made in paraffin oil (CAS No. 8012-95-1 Sigma-Aldrich), except for acetic acid, which was diluted in distilled water. A 1:100 dilution was used for the following odors: ethyl acetate, methyl butyrate, *i*-amyl acetate, prenyl acetate, methyl hexanoate, methyl benzoate, anisole, 1-hexanol, 2-hexanol, *E*-2-hexenal and g-octalactone. A 1:200 dilution was used for the following odors: methyl acetate, ethyl butyrate, butyl butyrate, 2- heptanone, citral, acetic acid, hexanoic acid, octanoic acid.

Larvae were removed from food vials by washing them out with a 15% sucrose solution. Supernatant larvae floating on the surface of the solution were selected with a fine brush, rinsed in distilled water, dabbed dry on tissue paper and then transferred to the Petri dish arena for behavioral tracking. Only larvae kept between 20 to 120 min after introduction of sucrose were used for behavioral experiments. The Petri dish was then sealed with its lid upon introduction of the larvae into the arena and loading of the odor onto the lid. The odor gradient was not pre-established before the introduction of the larvae. A group of ∼15 larvae was monitored in a single trial during 5 minutes for their chemotactic behavior in a controlled odor gradient. Larvae were thus exposed to an odor gradient in gaseous phase emanating from the two odor sources. The starting zone of the larvae was centered on the dish and the initial orientations of individual larvae effectively were random. Dishes were discarded after each trial. Larvae were used in single trials. The numbers of trials corresponding to the no-odor (paraffin-oil) control were: 28 for *D. melanogaster* and *D. yakuba*; 30 for *D. sechellia*, *D. erecta, D. suzukii* and *D. biarmipes*; 32 for *D. santomea*; 41 for *D. simulans*.

### Tracking and image processing

Larval behavior was tracked using a video camera (Scout Basler) and custom-made software written in LabView (National Instruments). Frames were acquired at 16 Hz and pre-processed in real-time online using the Multi-Worm Tracker (MWT) package (Swierczek, Giles et al. 2011), which recorded the posture skeletons and positions of individual larvae. The MWT also consists of the offline behavioral measurement software *Choreography*, which outputs time-series variables that describe larval movements and body postures for each individual larval trajectory. The data derived from *Choreography* was then analyzed in Matlab (MathWorks, Natick, MA) using custom-made programs to calculate additional time-series variables and to determine the distance of larvae to the source, preference index (see below), their reorientation behaviors and their locomotion speed.

### Data analysis and sensorimotor quantification

#### Preference index (PI)

Defined as the amount of time the larvae spent in the odor zones having a diameter of 2 cm (T_odorzone_) around both odor sources relative to the total amount of time tracked (T_total_). PI = T_odorzone_/T_total_

#### Distance to source

The distance of the larvae to the closest of the two odor sources and was computed by collecting the centroid positions of the larvae over time. To visualize how the distance varied with time, the centroid position of each larva to the closest odor source was collected over all time points, with data binned by intervals of 5 s. Data were pooled from all experiments to calculate an average value for each bin. A Savitzky-Golay smoothing filter was then applied of polynomial order one and frame length 9 (∼500 ms) to the averaged distance values.

#### Turns and turn rate

Turns are defined as events that involve the body displacement of the larva accompanied with a change in body orientation (heading) larger than 20°. Turns were identified as described in Schleyer, Reid et al. (2015) using the variables head angle, kink and curve identified using Choreography, and relying on changes in the larva’s reorientation speed (Gomez-Marin, Stephens et al. 2011) that had to pass a set of empirically derived Schmitt-trigger thresholds (Ohyama, Jovanic et al. 2013). The turn rate is defined as the sum of turns Turn_total_ divided by the duration of time larvae were tracked T_total_. Turn rate [turns/min] = Turn_total_/T_total_.

#### When-to-turn metric (Prop_turns while moving down-gradient_)

To visualize how the turn rate varies with the bearing of the larvae to the closest odor source, the turn rate was calculated over all bearing angles ([-180°, 180°], where 0° represents a bearing facing the source), as described in Schleyer, Reid et al. (2015). The turn rate toward the closest odor source (Turn rate_toward_) is calculated by counting all turn events during which larvae had an absolute bearing angle smaller than 90° (Turn_toward_) divided by the time during which larvae were tracked moving towards a source (T_toward_). Turn rate_toward_ = Turn_toward_/T_toward_. The turn rate away from a source (Turn rate_away_) includes all turn events during which larvae had an absolute bearing angle larger than 90° (Turn_away_) and is divided by the duration of time larvae were tracked moving away from a source (T_away_). Turn rate_away_ = Turn_away_/T_away_.

The *when-to-turn* metric was also quantified by calculating the proportion of turns that occurred while the animal was moving down the gradient (away from the source). This calculation offers a single value that could easily be visualized in a heat map (Figure 5Biii).

Prop_turns while moving down-gradient_ = Turn_while moving down-gradient_/Turn_total_.

#### Where-to-turn-to metric (Prop_turns directed toward_)

This metric quantifies the proportion of turns that were directed toward the closest odor source (Turn_directed toward_). It is calculated as the number of turns for which the absolute bearing angle after the turn was smaller than the absolute bearing angle before the turn, relative to the sum of turns (Turn_total_). Prop_turns directed toward_ = Turn_directed toward_/Turn_total_.

#### Run speed and run speed modulation index

Run speed is defined as the average locomotion speed [mm/s] of the larval midpoint (center of the spline) during runs. To visualize how the speed varies with the bearing of the larvae to the closest odor source, the speed was calculated over all bearing angles ([-180°, 180°], where 0° represents a bearing facing the source), with means calculated on bins of 10°. Data were pooled from all experiments to calculate an average value for each bin. The run speed modulation index was computed by integrating over the area under the curve taking the run speed at bearing (-180°) as the baseline of the curve. An approximate integral was obtained using the trapezoidal method (trapz function implemented in MATLAB). This integral provided us with a single value quantifying the extent of run speed modulation.

#### Weathervaning and weathervane index

This metrics quantifies the gradual reorientation a larva performs without making abrupt turning involving stopping. Following the same logic of Gomez- Marin and Louis (2014), the weathervane index calculates the change in body angle, defined as the angular change in the direction of the body axis, at each bearing angle to the source. To visualize how the body angle varied with the bearing angle to the closest odor source, the change in body angle was calculated over all bearing angles ([-180°, 180°], where 0° represents a bearing facing the source), with data binned by intervals of 20°. Data were pooled from all larvae per experiment to calculate an average body angle change value for each bin. The S-shape of the function follows a sinusoid shape; thus, the weathervane index was computed by weighting these binned averages with a sine function. The sum of mean body angle changes for left and right reorientations was then taken to obtain a single value quantifying the overall extent of *weathervaning*. The *weathervaning* index ranges between 0 (absence of correlation between the reorientation rate and the bearing) and 1 (perfect correlation between the reorientation rate and the bearing).

#### Physicochemical descriptors and principal components analysis (PCA)

We obtained the molecular 3D structures of each odorant from PubChem (https://pubchem.ncbi.nlm.nih.gov) and uploaded them into the commonly used and publicly accessible E-Dragon 1.0 program from VCCLAB (http://www.vcclab.org) which generated 1664 physicochemical molecular descriptors. We z-score normalized the descriptors before performing the PCA.

#### Hierarchical clustering

All dendrograms were built by performing a hierarchical clustering analysis on the behavioral data. Using the *pdist* Matlab function the similarity of dissimilarity between every pair of objects in our data set was measured by calculating the Euclidean distance between objects. Using the *linkage* Matlab function the objects were then grouped into a binary, hierarchical cluster tree by linking pairs of objects that are in close proximity based on the evaluated Euclidean distances. Objects thus become paired into binary clusters which are then grouped into larger clusters until a hierarchical tree is formed. By plotting the tree using the *dendrogram* Matlab function the natural partitioning of the data into groups can be visualized.

The definition of the three categories of odors in Figure 3B and Supplementary Fig. 2 is based on the partition of the data obtained through the hierarchical clustering analysis performed on the mean preference indices elicited by the different odors for each species.

#### Linear discriminant analysis

A default linear discriminant analysis classifier was created on the data using the *fitcdiscr* Matlab function. The coefficients for the linear boundary between the “turners” and “non-turners” was then retrieved and the curve separating them plotted. The behavioral data were then projected onto the axis perpendicular to the linear discriminant axis and to quantify the differences between the projected means of the “turners” and “non-turners”.

#### K-means clustering and silhouette value

The behavioral data was partitioned by means of the *k*-means clustering method. The squared Euclidean distance was used as distance metric to minimize the sum of the distances between the centroid and all member objects of the cluster. Each centroid is the mean of points in that cluster. Different *k*-means clustering solutions for different numbers of clusters were compared to determine an optimal number of clusters.

Cluster solutions were evaluated by examining silhouette plots and values created using the *silhouette* Matlab function (Supplementary Fig. 3B). To verify that our clustering solution was the optimal one, we also performed replicate clustering from different randomly selected centroids to ascertain that the solution with the lowest total sum of distances among all replicates was chosen.

## Statistical data analysis

The sample size of each (species - odor) condition of the full screen was determined based on the results of a preliminary screen where the Preference Index of the behavior of *D. melanogaster* was compared between a strong attractant Ethyl butyrate (1:100) and no odor. A sample size of 10 trials was found to be sufficient to ensure 98% of statistical power. Following the competition of the screen, this result was validated for each species by comparing the behavior of its HOP odor with the corresponding no-odor control. Statistical powers higher than 92% were found for all conditions. The statistical-power analysis was conducted by using the “sampsizepwr” function in Matlab (MathWorks, Natick, MA).

---

**Supplementary Figure 1:**
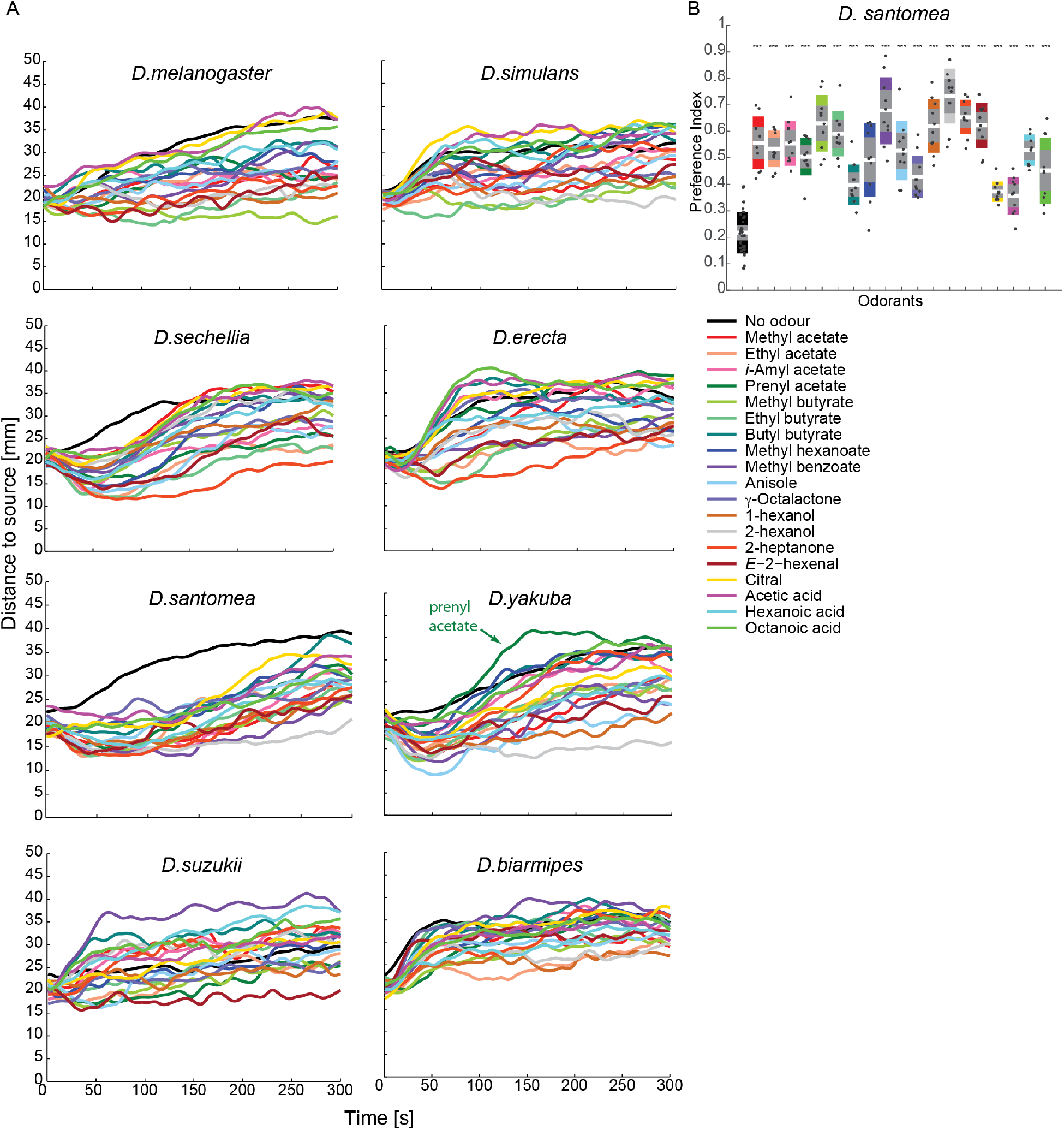
Time course of the preference index (PI). **(A)** Average distance of the larvae to the source over time for each species and each condition (see Materials and methods). **(B)** Boxplot of the PI calculated after 5 min for *D. santomea* tested with the 19 odors and the unstimulated condition. Each odor displays an average PI significantly higher than that of the unstimulated no-odor-control condition. Differences of mean of odor PIs with mean of no-odor PI (0.22) were tested by a one-sample one-sided *t*-Test (\**p*≤0.05, \*\**p*≤0.01, \*\*\**p*≤0.001, ns: not significant *p*>0.05). In each boxplot, the mean (white line) is surrounded by a gray area that corresponds to 1.96 SEM (95% confidence interval), and a colored interval that corresponds to 1 standard deviation (SD).

**Supplementary Figure 2:**
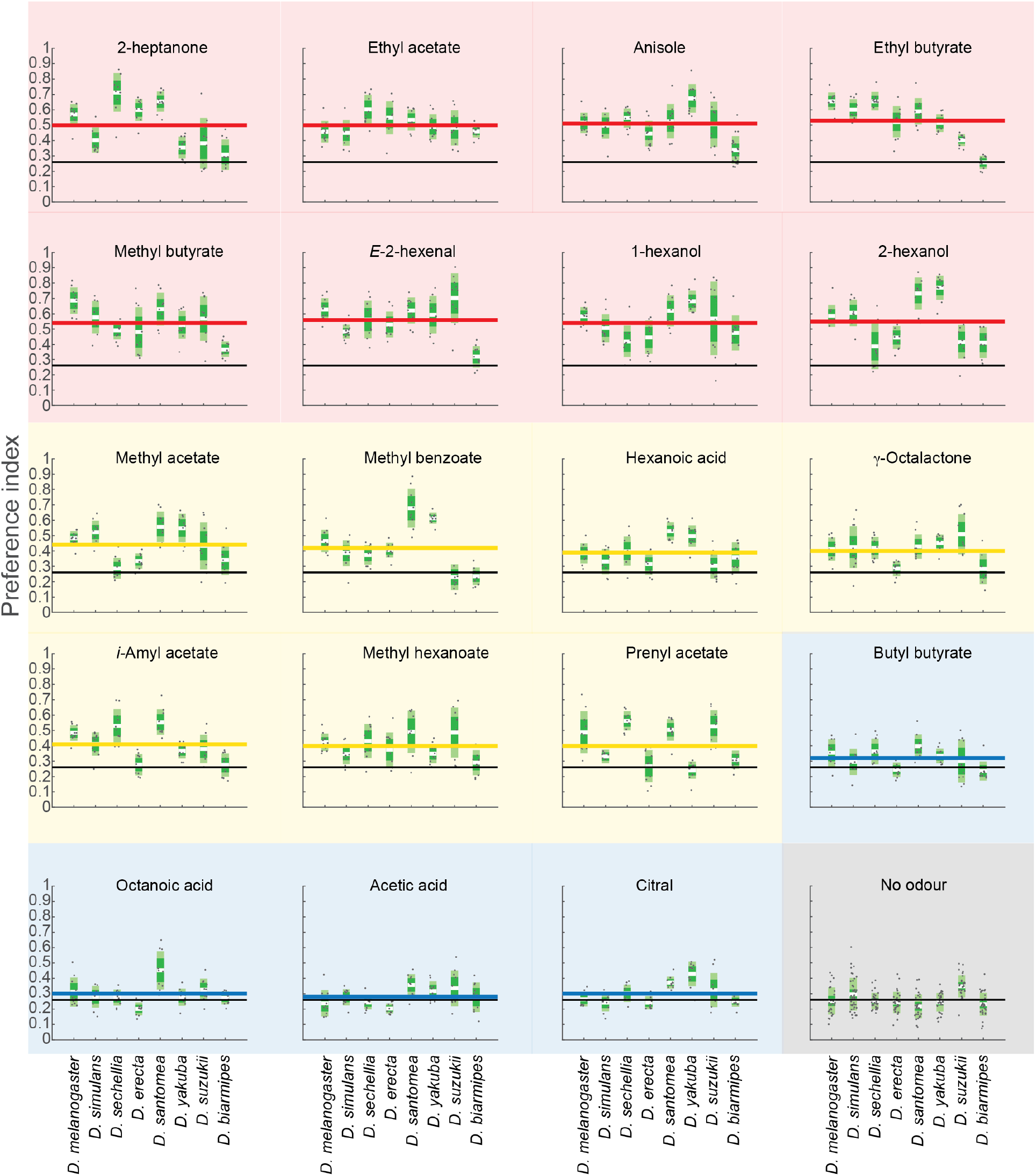
Boxplot of the preference index (PI) of each species for the 19 odors and the unstimulated condition as control. Every data point in a box plot is the average PI obtained from one trial (N=10 trials for the odor conditions and N=30 trials for unstimulated control condition). The white horizontal line in the plots represents the mean of all trials per condition. The mean (white line) is surrounded by a dark-green interval corresponding to 1.96 SEM (95% confidence interval) and light-green interval corresponding to 1 standard deviation. The long horizontal lines (red, yellow or blue) in each graph represent the average PI taken from the mean of each species for each condition. The black horizontal line is the average PI of all species for the unstimulated condition.

**Supplementary Figure 3:**
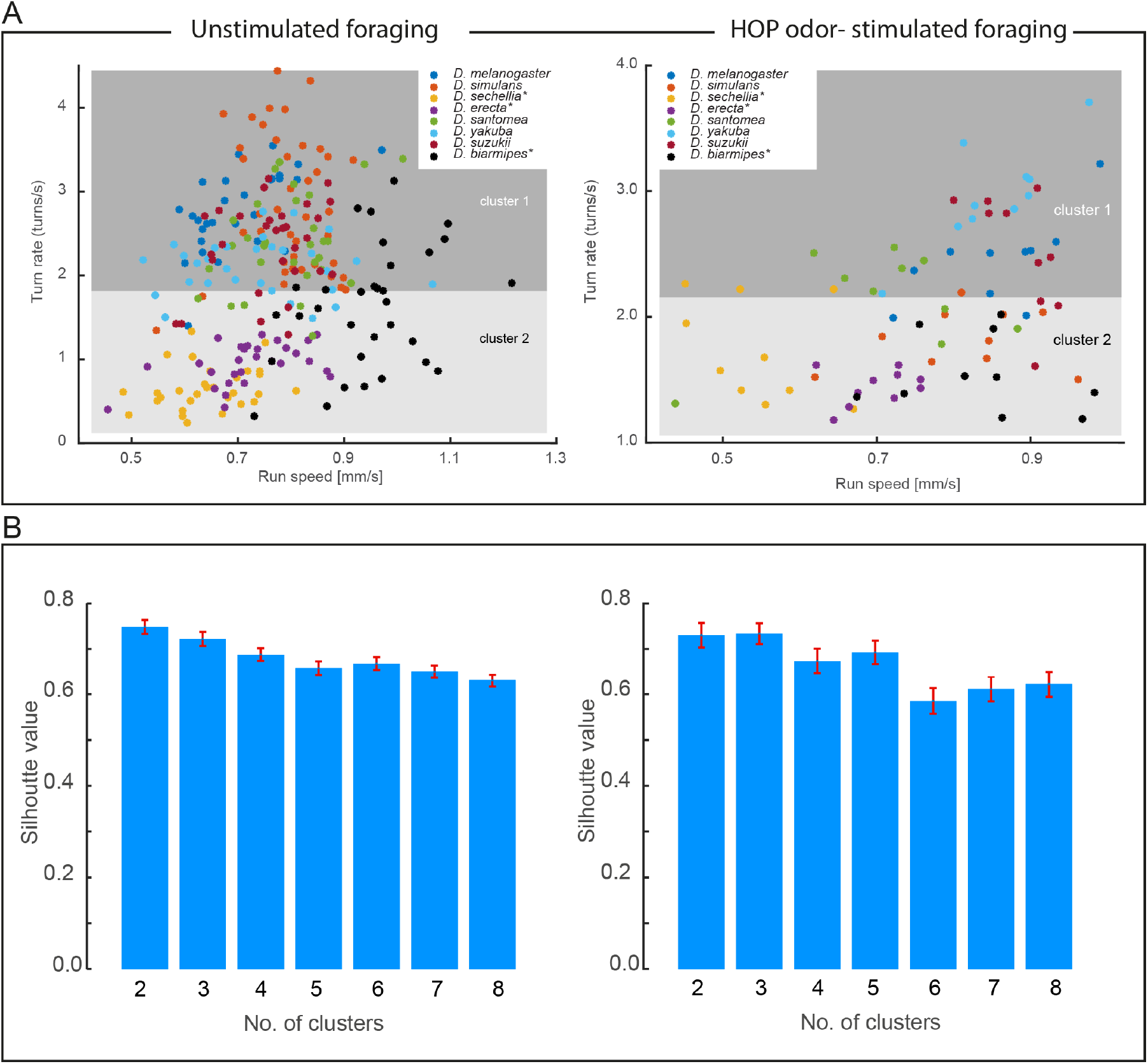
Clustering analysis of behavioral data. **(A)** *k*-means clustering was performed to partition the data into 2 clusters after which the clusters and cluster regions were plotted both for unstimulated (left) and HOP-stimulated (right) foraging. The names of the 3 species forming the non-turner group in Figure 4B are labeled with a star (*). The number of trials per species is 30 and 10 for the unstimulated and the HOP stimulated condition, respectively. **(B)** The silhouette value determines how well separated the clusters are. The quality of separation is highest for 2 and 3 clusters both for unstimulated and HOP-stimulated foraging. The error bars represent the SEM.

**Supplementary Figure 4:**
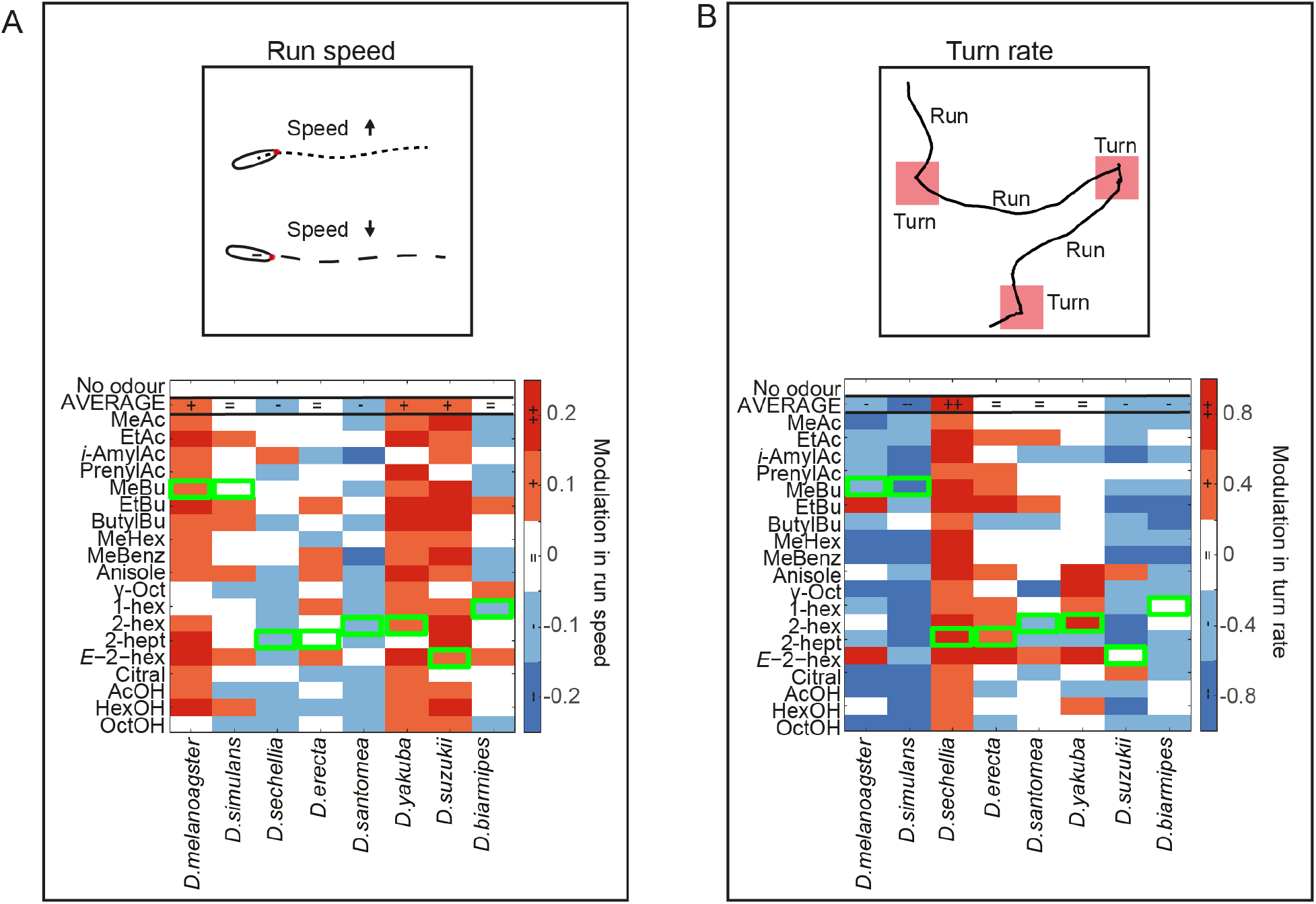
Analysis of changes in speed and turn rate upon odor stimulation. **(A)** Heat map of the run speed for all species and all tested odors relative to the no-odor condition of each species. The HOP odor of each species is demarked by an open green box. **(B)** Heat map of the turn rate for all species and all tested odors relative to the no-odor condition of each species. The HOP odor of each species is demarked by an open green box. Number of trials per experimental condition (species — odor): 10.

